# Comparative transcriptomic analysis reveals conserved transcriptional programs underpinning organogenesis and reproduction in land plants

**DOI:** 10.1101/2020.10.29.361501

**Authors:** Irene Julca, Camilla Ferrari, María Flores-Tornero, Sebastian Proost, Ann-Cathrin Lindner, Dieter Hackenberg, Lenka Steinbachová, Christos Michaelidis, Sónia Gomes Pereira, Chandra Shekhar Misra, Tomokazu Kawashima, Michael Borg, Frédéric Berger, Jacob Goldberg, Mark Johnson, David Honys, David Twell, Stefanie Sprunck, Thomas Dresselhaus, Jörg D. Becker, Marek Mutwil

## Abstract

The evolution of plant organs, including leaves, stems, roots, and flowers, mediated the explosive radiation of land plants, which shaped the biosphere and allowed the establishment of terrestrial animal life. Furthermore, the fertilization products of angiosperms, seeds serve as the basis for most of our food. The evolution of organs and immobile gametes required the coordinated acquisition of novel gene functions, the co-option of existing genes, and the development of novel regulatory programs. However, our knowledge of these events is limited, as no large-scale analyses of genomic and transcriptomic data have been performed for land plants. To remedy this, we have generated gene expression atlases for various organs and gametes of 10 plant species comprising bryophytes, vascular plants, gymnosperms, and flowering plants. Comparative analysis of the atlases identified hundreds of organ- and gamete-specific gene families and revealed that most of the specific transcriptomes are significantly conserved. Interestingly, the appearance of organ-specific gene families does not coincide with the corresponding organ’s appearance, suggesting that co-option of existing genes is the main mechanism for evolving new organs. In contrast to female gametes, male gametes showed a high number and conservation of specific genes, suggesting that male reproduction is highly specialized. The expression atlas capturing pollen development revealed numerous transcription factors and kinases essential for pollen biogenesis and function. To provide easy access to the expression atlases and these comparative analyses, we provide an online database, www.evorepro.plant.tools, that allows the exploration of expression profiles, organ-specific genes, phylogenetic trees, co-expression networks, and others.

## Introduction

The evolution of land plants has completely changed the appearance of our planet. In contrast to their algal relatives, land plants are characterized by three-dimensional growth and the development of complex and specialized organs. They possess a host of biochemical adaptations, including those necessary for tolerating desiccation and UV stress encountered on land, allowing them to colonize most terrestrial surfaces. The earliest land plants which arose ~470 million years ago ^1^, were speculatively similar to extant bryophytes, possessing tiny fertile axes or an axis terminated by a sporangium^2–4^. The innovation of shoots and leaves mediated the 10-fold expansion of vascular plants ^5,6^ and an 8–20-fold atmospheric CO2 drawdown ^7^, which significantly shaped the Earth’s geosphere and biosphere ^8^. To enable soil attachment and nutrient uptake, the first land plants only had rhizoids, filamentous structures homologous to root hairs ^9^. Roots evolved to provide increased anchorage (and thus increased height) and enable survival in more arid environments. Parallel with innovations of vegetative cell types, land plants evolved new reproductive structures such as spores, pollen, embryo sacs, and seeds together with the gradual reduction of the haploid phase. In contrast to algae, mosses, and ferns that require moist habitats, the male and female gametophytes of gymnosperms and angiosperms are strongly reduced, consisting of only a few cells, including the gametes ^10,11^. Moreover, sperm cells have lost their mobility and use pollen grains as a protective vehicle for long-distance transport and a pollen tube for their delivery deep into maternal reproductive tissues ^12,13^. The precise interaction of plant male and female gametes, leading to cell fusion, karyogamy, and development of both the embryo and endosperm after double fertilization has just begun to be deciphered at the molecular level ^14,15^. These anatomical innovations are mediated by coordinated changes in gene expression and the appearance of novel genes and/or repurposing of existing genetic material. Genes that are specifically expressed in these organs often play a major role in their establishment and function ^16,17^, but the identity and conservation of these specifically-expressed genes have not been extensively studied.

Nowadays, flowering plants comprise 90% of all land plants and serve as the basis for the terrestrial food chain, either directly or indirectly. The use of model plants like *Arabidopsis thaliana* and maize and technical advances allowing live-cell imaging of double fertilization have been instrumental for several major discoveries ^18,19^. When assessing current knowledge of male and female gamete development in plants, it is evident that the male germline has been studied to a greater extent ^11,20^. This is mainly due to its accessibility and the development of methods to separate the sperm cells from the surrounding vegetative cell of pollen, e.g. by FACS ^21^. Analysis of male germline differentiation, for example, has led to the identification of Arabidopsis *DUO POLLEN 1 (DUO1)* and the network of genes it controls, which include the fertilization factors, *HAP2/GCS1* and *GEX2*^22^. However, as novel genes are still being discovered that control the development of male and female gametes ^10,11^ or their functions ^23,24^, it is clear that our knowledge of the molecular basis of gamete formation and function is far from complete.

Current approaches to study evolution and gene function mainly use genomic data to reveal which gene families are gained, expanded, contracted, or lost. While invaluable, genomic approaches alone might not reveal the function of genes that show no sequence similarity to known genes^25^. To remedy this, we combined comparative genomic approaches with newly established, comprehensive gene expression atlases of two bryophytes *(Marchantia polymorpha, Physcomitrium patens),* a lycophyte *(Selaginella moellendorffii),* gymnosperms *(Ginkgo biloba, Picea abies),* a basal angiosperm *(Amborella trichopoda),* eudicots *(Arabidopsis thaliana, Solanum lycopersicum)* and monocots *(Oryza sativa, Zea mays).* We then compared these organ-, tissue- and cell-specific genes to identify novel and missing components involved in organogenesis and gamete development.

We show that transcriptomes of most organs are conserved across land plants and report the identity of hundreds of organ-specific gene families. We demonstrate that the age of gene families is positively correlated with organ-specific expression and the appearance of organ-specific gene families does not coincide with the appearance of the corresponding organ. We observed a high number of male-specific gene families and strong conservation of male-specific transcriptomes, while female-specific transcriptomes showed fewer specific gene families with less conservation. Our detailed analysis of gene expression data capturing the development of pollen revealed numerous transcription factors and kinases potentially important for pollen biogenesis and function. Finally, we present a user-friendly, online database www.evorepro.plant.tools, which allows the browsing and comparative analysis of the genomic and transcriptomic data derived from sporophytic and gametophytic samples across 13 members of the plant kingdom.

## Results

### Constructing gene expression atlases and identifying organ-specific genes

We constructed gene expression atlases for ten phylogenetically important species (Table 1). These include the bryophytes *Physcomitrium patens (Physcomitrella)* (Fig. 1a) and *Marchantia polymorpha* (Fig. 1b), the lycophyte *Selaginella moellendorffii,* the gymnosperms *Ginkgo biloba* and *Picea abies,* the basal angiosperm *Amborella trichopoda,* the monocots *Oryza sativa* and *Zea mays,* and the eudicots *Arabidopsis thaliana* and *Solanum lycopersicum* (Fig. 1c). The atlases were constructed by combining publicly available RNA sequencing (RNA-seq) data with 134 fastq files generated by the EVOREPRO consortium (see Supplementary Table 1). For each species, we generated an expression matrix that contains transcript-level abundances captured by transcript per million (TPM) values ^26^. The expression matrices capture gene expression values from the main anatomical sample types, which we grouped into ten classes: flower, female, male, seeds, spore, leaf, stem, apical meristem, root meristem, and root (Fig. 1a-c). Furthermore, the expression data was used to construct co-expression networks and to create an online EVOREPRO database allowing further analysis of the data (www.evorepro.plant.tools).

**Table 1.** Organs, tissues, and cell types used in the expression atlases analyzed. The different species are shown in columns, while the rows organize the organs, tissues and cell types into rows.To summarize, these results show that organ-specific genes represent a significant part of the transcriptome, with male and root samples possessing the most specialized transcriptomes.

| Organ/tissue/cell type | Marchantia | Physcomitrella | Selaginella | Ginkgo | Spruce | Amborella | Arabidopsis | Tomato | Rice | Maize |
| --- | --- | --- | --- | --- | --- | --- | --- | --- | --- | --- |
| Flower | N/A | N/A | Strobili | Strobili, microstrobilus | N/A | Tepals, buds, opened flowers | Buds, stamen filaments, carpels, petals, stigmatic tissue, sepals | Buds, opened flowers | Buds, panicles | Tassels, ear, silk |
| Apical meristem | - | - | - | - | - | Apical meristem | Apical meristem | Apical meristem | Apical meristem | Apical meristem |
| Male | Sperm | Sperm | - | - | - | Pollen (mature, tube), sperm, microspores, generative cell | Pollen (mature, tube, bicellular, tricellular), microspore, sperm | Pollen (mature, tube), microspore, sperm cell, generative cell | Pollen (bi-, tricellular), microspore, sperm | Pollen (mature, tube), microspore, sperm |
| Female | - | - | - | Ovules | - | Ovary, egg apparatus cell | Egg cell, ovule | Ovaries, ovary walls, ovules | Ovary, ovule, egg cell | Ovary, ovule, nucellus, egg cell, embryo sac |
| Root | N/A | N/A | Roots, rhizophores | Roots | - | Roots | Root (apex, tip, differentiation zone, stele, elongation zone) | Root (differentiation zone, elongation zone) | Root (differentiation zone, elongation zone) | Root (tip, secondary, stele, elongation zone, maturation zone) |
| Root meristem | N/A | N/A | Meristematic zone | - | - | - | Meristematic and QC zone | Meristematic zone | Meristematic zone | Root (meristematic zone) |
| Leaf | Thallus | Leaflets | Microphyll | Leafs | Needles | Leaves | Leaves | Leaves | Leaves | Leaves |
| Stem | N/A | N/A | Top stem, bottom stem | Stem, xylem, cambium | Xylem, phloem, cambium | - | Stems | Stems | Stems | Stems |
| Seed | N/A | N/A | N/A | Kernels | - | - | Seeds (young, germinating), endosperm | Seeds (5-30 DPA) | Seeds | Seeds (mature, germinating), endosperm,<br>pericarp and<br>aleurone |
| Spore | Sporeling | Spore<br>capsule,<br>germinati<br>ng spores | - | N/A | N/A | N/A | N/A | N/A | N/A | N/A |

**Fig. 1:**
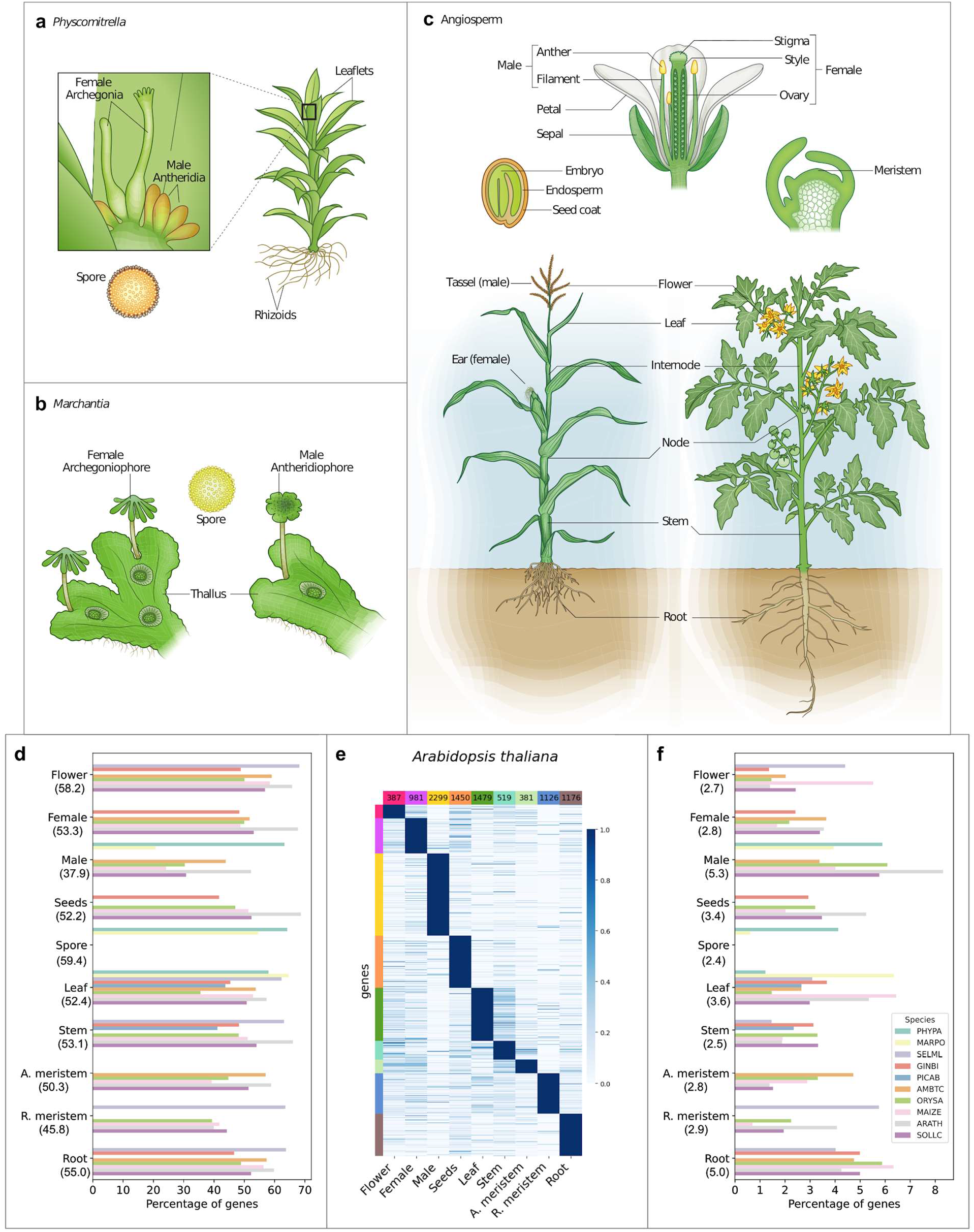
Expression atlases for seven land plant species. Depiction of the different organs, tissues, and cells collected for (**a**) *P. patens* (**b**) *Marchantia polymorpha,* and (**c**) angiosperms. **d**, The percentage of genes (x-axis) found to be expressed (defined as TPM>2) in organs (y-axis) of the different species (indicated by colored bars as in (f)). The numbers beneath the organs (y-axis) indicate the average percentage of genes for all species. **e**, Expression profiles of organ-specific genes from *Arabidopsis thaliana.* Genes are in rows, organs in columns and the genes are sorted according to the expression profiles (e.g., flower, female). The numbers at the top of each column indicate the total number of genes per organ. Gene expression is scaled to range from 0-1. Bars on the left of each heatmap show the sample-specific genes and correspond to the samples on the bottom: pink - Flower, purple - Female, yellow - Male, orange - Seeds/Spore, dark-green - Leaf, medium-green - Stem, light-green - Apical meristem, blue - Root meristem, brown - Root. **f**, The percentage of genes with specific expression in the ten species.

To identify genes expressed in the different samples, we included only those with an average TPM >2 (see methods). For all ten species, approximately 71% of their genes were expressed in at least one structure (Supplementary Table 2). Interestingly, the male sample has a lower percentage (38%) followed by root meristems (46%), while the other samples have between 50-60% expressed genes (Fig. 1d).

Organ- and cell-specific genes can often play a major role in the establishment and function of the organ and cell type ^16,17^. To identify such genes, we calculated the specificity measure (SPM) of each gene, which ranges from 0 (not expressed in a sample) to 1 (expressed only in the sample). A threshold capturing top 5% of the SPM values was used to identify the sample-specific genes for all species (Supplementary Fig. 1, Supplementary Table 3). To examine the sample-specific genes’ expression profiles, we plotted the scaled TPM values of these genes for *A. thaliana.* Visual inspection shows that the TPM values of the samplespecific genes are in all cases highest in the samples that the genes are specific to (Fig. 1d, Supplementary Fig. 2). For the ten species, an average of 21% of the genes were identified as sample-specific (Supplementary Table 2). The lowest percentage was found in *P. abies* (5%), followed by *M. polymorpha* (11%) and *P. patens* (11%), while the highest percentage was found in *A. thaliana,* where 35% of the transcripts showed sample-specific expression (Supplementary Table 2). These low and high percentages observed can be partially explained by the number of organs and cell types that we analyzed (Supplementary Table 1).

Interestingly, we observed that the male (5.3%) and root (5.0%) samples typically contained the highest percentage of specific genes (Fig. 1f, Supplementary Table 2). In *A. thaliana,* the higher percentage of male-specific genes was in agreement with previous studies that showed a high specialization of the male transcriptome ^27,28^. Conversely, stem, spore, apical meristem, root meristem, flower, and female show values lower than 3% (Fig. 1f, Supplementary Table 2). Previous studies also showed the low number of genes mainly expressed in the female gametophyte ^29,30^.

### Are the transcriptomes of organs conserved across species?

Our above analysis suggests that sample-specific gene expression is widespread, and we set out to investigate whether these patterns are conserved across species. To this end, we investigated which samples specifically expressed similar sets of gene families (represented by orthogroups) by employing a Jaccard distance that ranges from 0 (two samples express an identical set of sample-specific gene families) to 1 (none of the sample-specific gene families are the same in the two samples). We expected that if, e.g., the root-specific transcriptome is conserved across angiosperms, then Jaccard distance of root vs. root transcriptomes (e.g., *Arabidopsis* root vs. rice root) should be lower than when comparing root vs. non-root transcriptomes (e.g., *Arabidopsis* root vs. rice leaf).

The analysis revealed that *Arabidopsis* flower-, male-, seeds-, stem- and root-specific transcriptomes were significantly more similar to the corresponding sample in the other species (p-value < 0.05, Fig. 2a). When performing the analysis for all ten species, we observed that root, male, and seeds expressed specifically similar gene families in all species with the samples (7 species for root, 7 for male, and 5 for seeds) and for other organs, some species show significance, flowers (5 out of 7 species with flower samples), female (2 out of 6), leaf (7 out of 10), stem (5 out of 7), apical meristem (4 out of 5), root meristem (4 out of 5) (Fig. 2b, Supplementary Fig. 3). Conversely, spore (0 out of 2) samples did not show similar transcriptomes across *Marchantia* and *Physcomitrella* (Fig. 2b, Supplementary Fig. 3). We also performed clustering analysis between all pairs of sample-specific genes in the ten species and observed root-, seed-, flower, leaf-, meristem- and male-specific clusters (Supplementary Fig. 4). Interestingly, the male samples in *Physcomitrella* and *Marchantia* formed a distinctive cluster (Supplementary Fig. 4), suggesting that flagellated sperm of bryophytes employ a unique male transcriptional program compared with non-motile sperm of angiosperms.

**Fig. 2:**
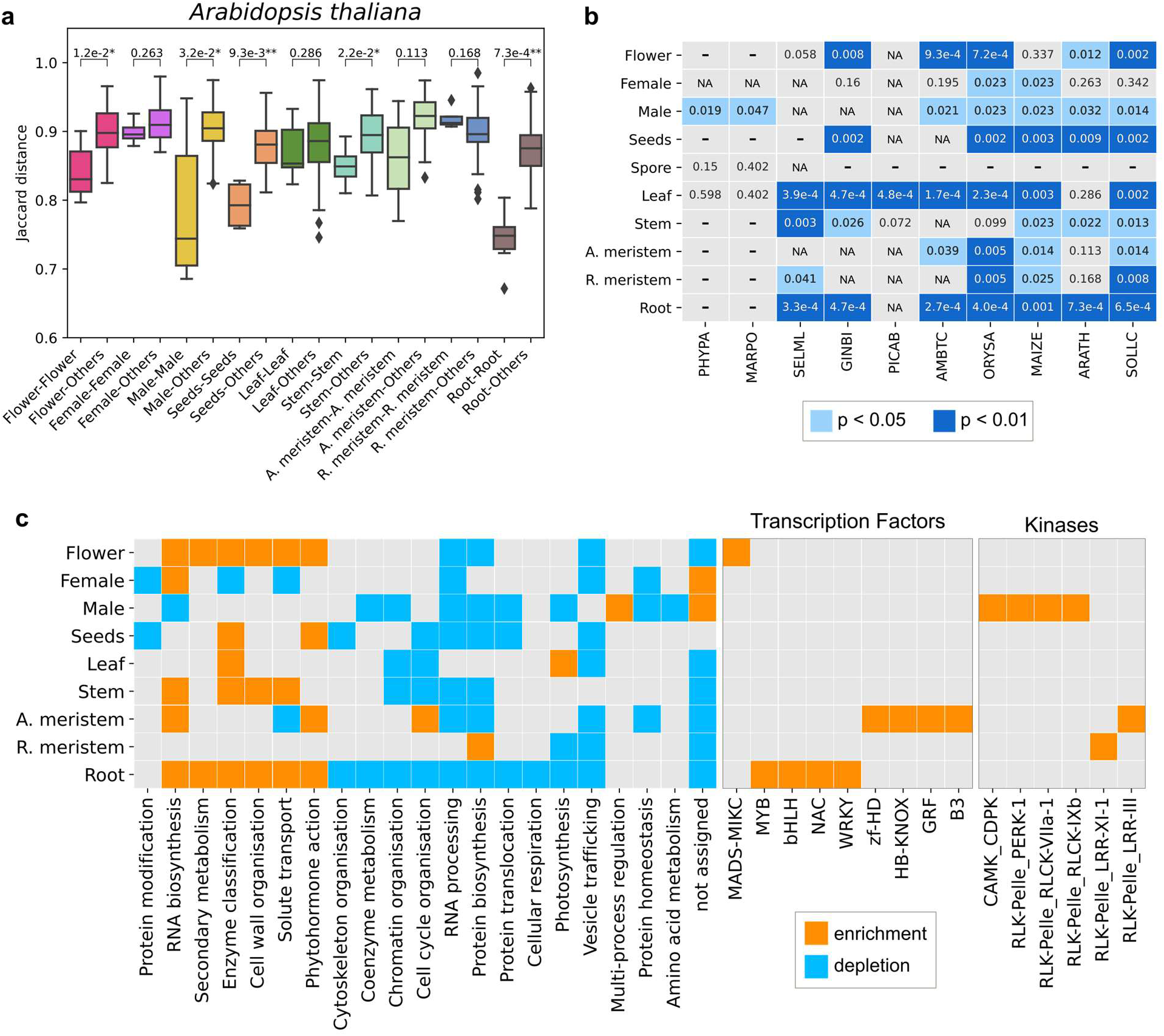
Comparison of sample-specific transcriptomes. **a**, Bar plot showing the Jaccard distances (y-axis) when comparing the same samples (x-axis, e.g., male-male) and one sample versus the others (e.g., male-others) for *Arabidopsis thaliana.* Lower values indicate a higher similarity of the transcriptomes. **b**, Significantly similar transcriptomes are indicated by blue cells (light blue p<0.05 and dark blue p<0.01). Species are indicated by the mnemonic: PHYPA - *P. patens,* MARPO - *Marchantia polymorpha,* SELML - *Selaginella moellendorffii,* GINBI - *Ginkgo biloba,* PICAB - *Picea abies,* AMBTC - *Amborella trichopoda,* ORYSA - *Oryza sativa,* MAIZE - *Zea mays,* ARATH - *Arabidopsis thaliana,* SOLLC - *Solanum lycopersicum.* **c**, Heatmap showing the significant (p-value < 0.05) functional enrichment (orange cell) or depletion (blue cell) in the ten sample classes (y-axis) in at least 50% species. The heatmap indicates Mapman bins (photosynthesis-not assigned), transcription factors, and kinases.

To reveal which biological processes are preferentially expressed in the different samples across the ten species, we performed a functional enrichment analysis of Mapman bins, transcription factors, and kinases (Fig. 2c, Supplementary Fig. 5). The analysis revealed that many functions were depleted in male and root samples in at least 50% of the species, indicating that most male and roots’ cellular processes were significantly repressed (p-value < 0.05, Fig. 2c, Supplementary Fig. 5). As expected, genes associated with photosynthesis were enriched in leaves but depleted in roots, root meristems, and male samples. Genes expressed in roots were enriched in solute transport functions, enzyme classification (enzymes not associated with other processes), RNA biosynthesis, secondary metabolism, phytohormone action, and cell wall organization (Fig. 2c). Interestingly, female and male reproductive cells were enriched for ‘not assigned’ bin, indicating that these organs are enriched for genes with unknown functions.

Since the sample-specific genes (Supplementary Table 3) are likely important for the formation and function of the organ, we investigated sample-specific transcription factors (Supplementary Table 4) and receptor kinases (Supplementary Table 5). An enrichment analysis of transcription factors (69 families) and kinases (142 families) showed that apical meristem and root samples were highly enriched in transcription factors, while male and apical meristem were enriched for kinases (Fig. 2c). In apical meristems, some of the enriched transcription factor families (C2C2-YABBY, GRF) were associated with the regulation, development, and differentiation of meristem ^31,32^. In roots, the enriched transcription factors (MYB, bHLH, WRKY, NAC) are related to biotic and abiotic stress response and root development ^33–37^. These sample-specific genes are thus prime candidates for further functional analysis (Supplementary Table 5).

### Phylostratigraphic analysis of sample-specific gene families

Organs, such as seeds and flowers, appeared at a specific time in plant evolution. To investigate whether there is a link between gene families’ appearance and their expression patterns, we used the proteomes of 23 phylogenetically important species and a derived species tree based on One Thousand Plant Transcriptomes Initiative (2019). Each orthogroup was placed to one node (phylostrata) of the species tree, where node 1 indicated the oldest phylostratum, and node 23 indicated the youngest, species-specific phylostratum (Supplementary Table 6). A total of 131,623 orthogroups were identified in the 23 Archaeplastida, of which 113,315 (86%) were species-specific, and the remaining 18,308 (14%) were assigned to internal nodes. Of these internal node orthogroups, most were ancestral (24% - node 1, 10% - node 3), belonged to streptophytes (7%, node 6), land plants (7%, node 8), seed plants (10%, node 13), monocots (0.3%, node 18), or eudicots (1%, node 19) (Fig. 3a). Analysis of phylostrata in each species revealed a similar distribution of the orthogroups, with most of them belonging to node 1 (~34%) or being species-specific (~31%, Supplementary Fig. 6).

**Fig. 3:**
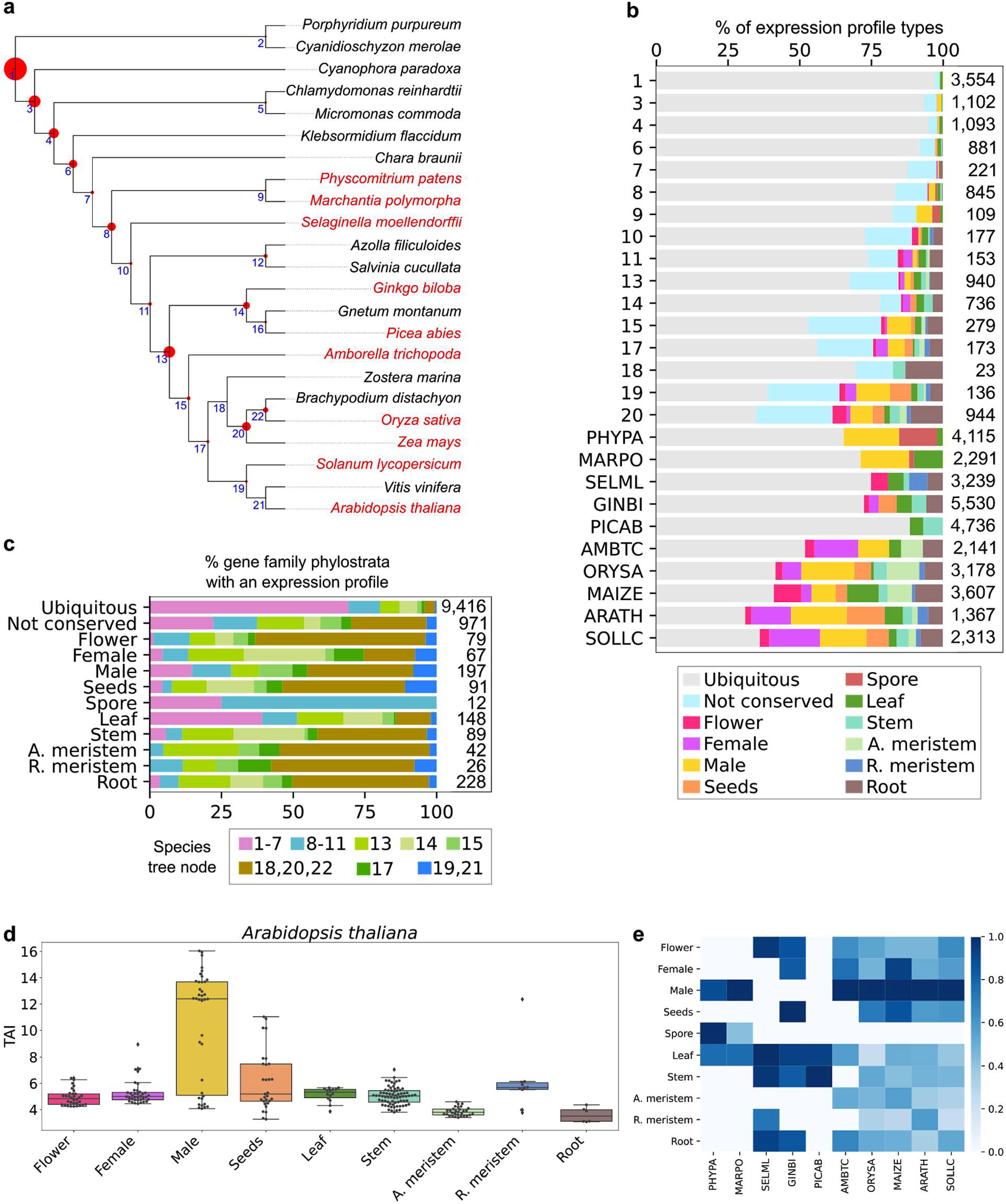
Genomic analysis of sample-specificity of gene families. **a**, Species tree of the 23 species for which we have inferred orthogroups. Species in red are the ones with transcriptomic data available. Blue numbers in the nodes indicate the node number (e.g., 1: node 1). The tree’s red circles show the percentage of orthogroups found at each node (largest: node 1 - 24% of all orthogroups, smallest: node 21 - 0.1%). **b**, Percentage of expression profile types of orthogroups per node. The expression profile types are: ubiquitous (light gray, orthogroup is not organ-specific), not conserved (light blue, organ-specificity not conserved in different species), or sample-specific (e.g., brown: root-specific). **c**, Percentage of phylostrata (nodes) within the different expression profile types. **d**, Transcriptome age index (TAI) of the different sample-specific genes in *Arabidopsis thaliana.* The boxplots show the TAI values (y-axis) in the different organs (x-axis), where a high TAI value indicates that the sample expresses a high number of younger genes. **e**, Summary of the average TAI value in the ten species. The organs are shown in rows, while the species are shown in columns. The TAI values were scaled to 1 for each species by dividing values in a column with the highest column value.

To investigate whether the different phylostrata show different expression trends, we surveyed orthogroups that contain at least two species with RNA-seq data, which resulted in 37,887 (29% of the total number of orthogroups) meeting this criterion. Then, each orthogroup was assigned to different expression profiles: ubiquitous (not specific in any organ), not conserved (e.g., root-specific in one species, flower-specific in others), or organ-specific (for details see material and methods, Supplementary Table 6). The majority of the orthogroups in internal nodes (not species-specific) of the phylogenetic tree were assigned as ubiquitous (9,416), which corresponded to orthogroups that showed broad and not organ-specific expression (Fig. 3b). Interestingly, we observed a clear pattern of gene families becoming increasingly organ-specific as phylostratigraphic age decreased (<5% specific genes in node 1, vs. ~25% in node 13), indicating that younger gene families are recruited to specific organs (Fig. 3b).

Next, we identified sample-specific gene families and investigated when they appeared during plant evolution. The number of gene families in internal nodes per sample varied from 12 (spore) to 228 (root), and we observed trends of samples across the internal nodes. In general, many organ-specific orthogroups were present in nodes corresponding to monocots (Node 18, 20, 22). Expectedly, the 9,416 ubiquitous orthogroups were mostly of ancient (node 1-7) origin, suggesting that these old gene families tend to show a broader expression. The nonconserved groups had both old and more recent gene families. From the organ-specific families, leaves and spores were the groups containing more ancient families, while meristems had younger families. Flower, root, seeds, stem had few older families. Interestingly, when we compared male and female groups, we observed that the male-specific orthogroups had older gene families than the female-specific orthogroups (Fig. 3c).

Several studies revealed that new genes in animals tend to be preferentially expressed in male reproductive tissues, such as testis ^38–40^. Similar observations have been made in Arabidopsis, rice, and soybean ^41^, where new genes were predominantly expressed in male reproductive cells ^42^, suggesting that these cells may act as an “innovation incubator” for the birth of *de novo* genes. Our gene expression data also revealed that male samples possess the youngest transcriptome in Arabidopsis (Fig. 3d, yellow bar), and in the male samples of *M. polymorpha, A. trichopoda, Z. mays, O. sativa, S. lycopersicum,* but not in *P. patens* (Fig. 3e, dark-blue cells for male, Supplementary Fig. 7). With the unclear exception in *Physcomitrella,* we conclude that the observation that male samples express young genes is robust in the plant kingdom. However, pollen also expresses a substantial portion of old genes (species nodes 1-7 in Figure 3c), probably representing an old transcription program present in gametes in Archaeplastida.

### Phylostratigraphic and gene expression analysis reveals that co-option drives the evolution of organs

The evolution of land plants involved many major innovations mediated by gains and losses of gene families and co-option of existing gene functions. Most of the changes are related to land adaptations comprising requirements for structural support, uptake of water, prevention of desiccation and gas exchange^43^. To better understand this complex process, we first analyzed the enrichment/depletion of organ-specific and ubiquitous genes in each node of the species tree (Supplementary Table 7). In line with previous results (Figure 3b), ubiquitous genes were enriched for genes that appeared before the divergence of land plants and depleted for genes that appeared when plants colonized land (node 8, Fig. 4a). In line with the basal function (photosynthesis) of leaves, leaf-specific genes were enriched in ancestral nodes and the species-specific nodes of *M. polymorpha* (thallus samples) and *S. moellendorffii* (microphyll), and depleted in species-specific nodes of the seed plants (Fig. 4a).

**Fig. 4:**
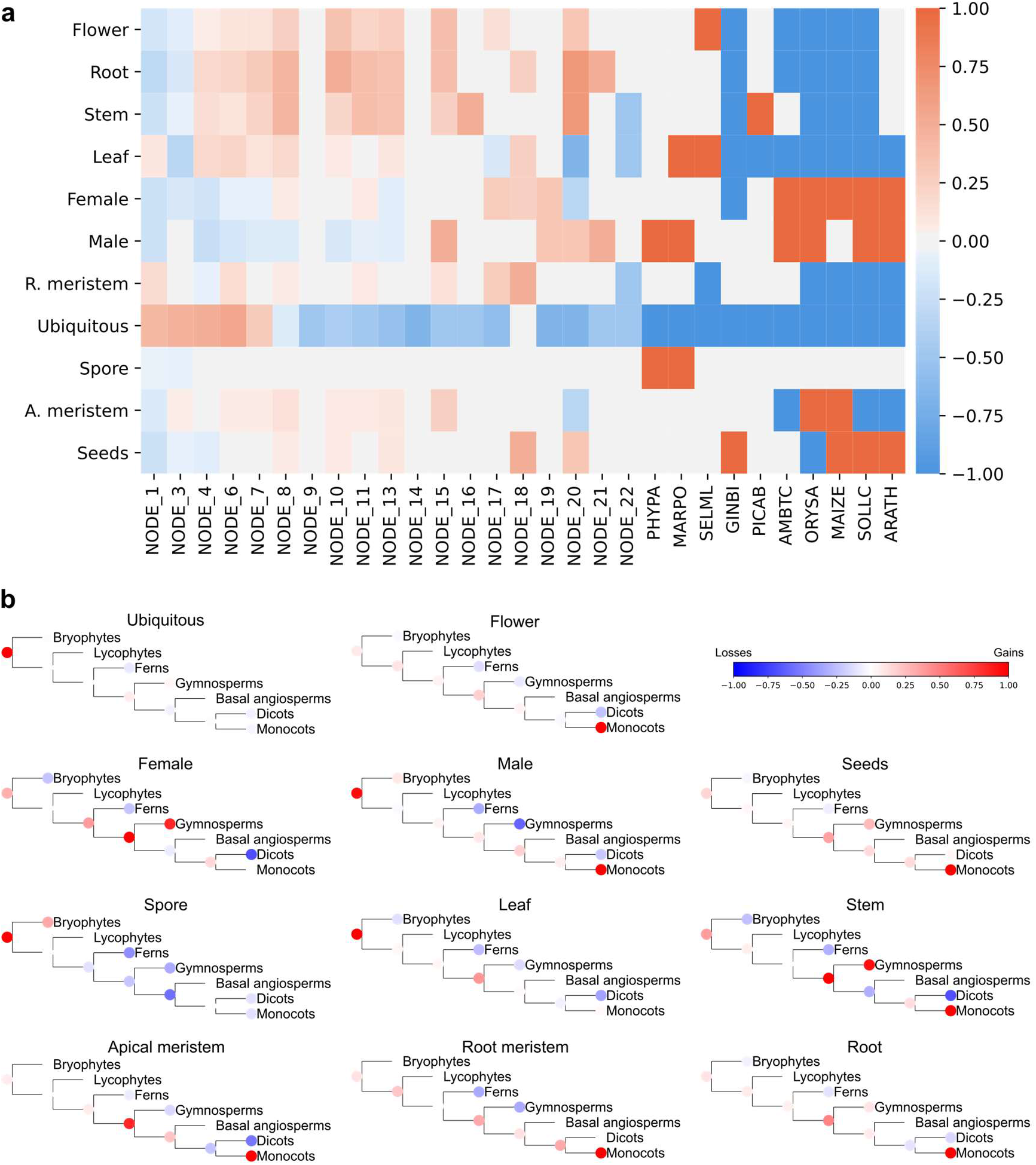
Evolutionary analysis of organs. **a**, Enrichment and depletion of organ-specific genes per node in the species tree (nodes are the same as in Fig. 3a). The colors correspond with the number of species showing enrichment in each case (dark red: all species show enrichment, dark blue: all species show depletion). **b**, Cladograms of the main lineages showing gain (in red) and loss (blue) of gene families with ubiquitous and sample-specific expression profiles.

Leaf-specific gene families were acquired mainly in two ancestral nodes, before the divergence of land plants and before the divergence of seed plants (Fig. 4b). Most of the gene families were gained in node 1 (34 families, Supplementary Table 8). Leaves have multiple origins in land plants ^44,45^, however, the programs for oxygenic photosynthesis originated in ancient organisms ^46^. In agreement, before the divergence of land plants, we observed enrichment for functions related to photosynthesis (<N8), and after the divergence of land plants, we detected enrichment for additional functions such as external stimuli response, cytoskeleton organization, phytohormone action, and protein modification (Supplementary Table 9).

Interestingly, stem-, root-, and flower-specific genes shared a similar pattern and appeared to be enriched in nodes 4-8, 10-13, 15, and 20, and depleted in the species-specific nodes of vascular plants, except for *P. abies* for stems and *S. moellendorffii* for flowers. Although the origins(s) of roots, stems, and flowers are associated with vascular plants ^47–49^, we observed gene family expansions before the divergence of land plants (Fig. 4b) and in nodes as old as node 3 (2 orthogroups) for stems, node 1 (1 orthogroup) for roots, and node 3 (1 orthogroup) for flowers (Supplementary Table 8). Previous studies suggested that the evolution of novel morphologies was mainly driven by the reassembly and reuse of pre-existing genetic mechanisms ^45,50^. It was indicated that primitive root programs may have been present before the divergence of lycophytes and euphyllophytes ^51^. Also, before the divergence of charophytes from land plants, an ancestral origin was proposed for the SVP subfamily, which plays a crucial role in the control of flower development ^52,53^. A recent study has shown that a moss (*Polytrichum commune*) possesses a vascular system functionally comparable to that of vascular plants ^54^. These results support the idea that primitive stem-, root-, and flower-specific gene families existed prior to vascular plants’ divergence. After the divergence of land plants, we can observe that there is incremental gene family gain in monocots for all three organs (roots, stems, flowers, Fig. 4b, indicated by red nodes), and also to a lesser extent in the ancestral node of seed plants. Specifically, for stem, we observed more gains in gymnosperms and more losses in eudicots. Functional enrichment analysis supports only enrichment in nodes corresponding to land plants (>N8) and not in older nodes (Supplementary Table 9).

Apical and root meristem-specific genes appeared enriched in ancestral nodes and depleted in species-specific nodes, with the exception of apical meristem in monocots that are enriched (Fig. 4a). The analysis of gain/loss of gene families showed that many apical meristem-specific orthogroups were gained in seed plants and monocots and lost in eudicots. For root meristem-specific gene families we observed that many orthogroups were gained in monocots (Fig. 4b). Functional enrichment analysis for apical meristem-specific gene families shows enrichment of unknown functions in nodes N19 and N20, and for root meristem-specific gene families shows enrichment for phytohormone action in N8 and protein modification in N15 (Supplementary Table 9).

Seed-specific genes were enriched only in nodes of land plants. The nodes that showed enrichment were N10 (vascular plants), N18 and N20 (monocots), and species-specific nodes with the exception of *O. sativa,* which showed depletion of this set of genes. Some seed-specific families were gained before the divergence of land plants, but interestingly the higher number of gains was observed in N20 (monocots - 39 gene families), followed by N14 (gymnosperms - 15), N13 (seed plants - 11), and N19 (eudicots - 10) (see Fig. 4b, Supplementary Table 8). Enrichment of functions related to solute transport was observed only in eudicots (N19, Supplementary Table 9).

Spore-specific genes were enriched only in the species-specific nodes of bryophytes (Fig. 4a). However, gene family gains were observed in ancestral nodes (N4, N6, N8, N9, see Supplementary Table 8) and lipid metabolism enrichment only in the node ancestral to bryophytes (N9, Supplementary Table 9).

Male-specific genes were enriched in angiosperms (N15), monocots (N20), eudicots (N19, N21), and species-specific nodes, while female-specific genes were enriched only in monocots (N18, N22), eudicots (N19), and species-specific nodes (Fig. 4a). Additional male-specific families were gained in older nodes than female-specific families (intensity of the red color in the ancestral node of land plants, Fig. 4b). For male gene families, we observed six waves of gains (>15 gene families) in nodes N3, N8 (land plants), N13 (seed plants), N15 (angiosperms), N19 (eudicots), N20 (monocots). From these nodes, parallel to gains, we also observed many losses (>=10 gene families) in three nodes N13 (seed plants), N15 (angiosperms), and N19 (eudicots) (Supplementary Table 8). For female-specific families, we observed three main waves of gains (>10 gene families) in nodes N13 (seed plants), N14 (gymnosperms), N20 (monocots), and different waves of losses (Supplementary Table 8). Male gene families showed enrichment for protein modification, enzyme classification, RNA biosynthesis, cell cycle organization, phytohormone action, and female gene families showed enrichment only for RNA biosynthesis (Supplementary Table 9). Considering gains and losses of gene families, male-specific families were gained mainly in the node ancestral to land plants, and in monocots, and for female-specific families in seed plants and gymnosperms (Fig. 4b).

In summary, the genetic programs for organ-specific genes are present in older nodes, before the divergence of land plants. Monocots seem to be the group with more gene family gains, which is in agreement with previous studies ^55^.

### Comparisons of transcriptional programs of gametes

Sexual reproduction is a complex process. In diploid flowering plants involves the production of haploid male and female gametes and fertilization of the female ovule by male gametes mediated by pollination (Fig. 5a). The pollen delivers the sperm cell(s) to the ovary by a pollen tube, and the fertilized ovules grow into seeds within a fruit (Fig. 5a). The two haploid bryophytes in our study differ in their sexual reproduction. *Physcomitrella* is monoicous and bears both sperm and eggs on one individual (Fig. 5b), and *Marchantia* is dioicous and bears only egg or sperm, but never both (Fig. 5c). However, both species produce motile sperm that require water droplets to fertilize the egg, generating diploid zygotes. The zygotes divide by mitosis and grow into a diploid sporophyte. The sporophyte eventually produces specialized cells that undergo meiosis and produce haploid spores, which are released and germinate to produce haploid gametophytes (Fig. 5b,c).

**Fig. 5:**
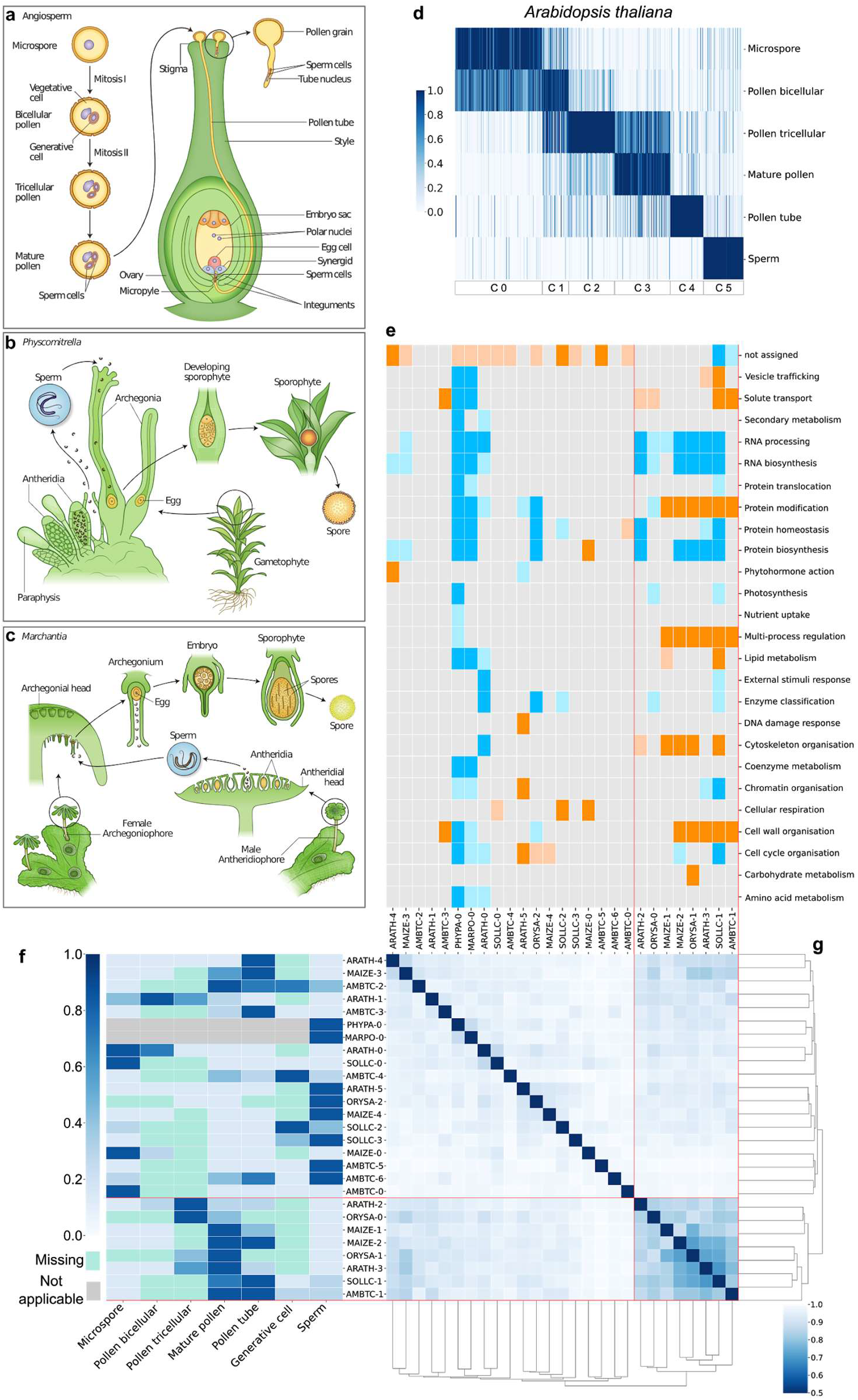
Comparison of male development across species. Overview of sexual reproduction in (**a**) Angiosperms, (**b**) *Phycomitrella,* and (**c**) *Marchantia.* **d**, Heatmaps showing the expression of male samples genes for *Arabidopsis thaliana.* Genes are in columns, sample names in rows. Gene expression is scaled to range between 0-1. Darker color corresponds to stronger gene expression. Bars to the bottom indicate the k-means clusters. **e**, Heatmap showing enrichment (orange) and depletion (blue) of functions in the found clusters. Light colors: p<0.05, dark colors: p< 0.01. **f**, Heatmap showing the average normalized TPM value per cluster for all the species. **g**, Clustermap is showing the Jaccard distance between pairs of clusters of all the species.

To further study whether the transcriptional programs of sexual reproduction are conserved in land plants, we applied k-means clustering on the male- and female-specific genes over the RNA-seq samples representing different samples of male and female organs (Supplementary Table 1). For male-specific genes, the analysis assigned each sample to one or more clusters (Fig. 5d exemplifies male samples in Arabidopsis (for other species, see Supplementary Fig. 8), with a variable number of genes assigned to each cluster (Supplementary Table 10). We then inferred which biological processes were enriched in the clusters (Fig. 5e), plotted an average expression profile of the genes in each cluster (Fig. 5f), and used Jaccard distance to identify similar clusters across species (Fig. 5g). Interestingly, three clusters showed strong similarity and were specific to pollen tricellular, mature pollen, and pollen tube for Angiosperms (Fig. 5g, indicated by red lines). Functional enrichment analysis revealed that pollen tricellular, mature pollen, and pollen tube samples were mainly enriched for cell wall organization, cytoskeletal organization, multi-process regulation, and protein modification (supported by five species, Fig. 5e). Conversely, other clusters showed enrichment for genes without assigned functions, and depletion for many biological processes (Fig. 5e).

Female samples included were less diverse than male samples. In all species, each sample was assigned to a cluster with exception of *O. sativa,* where ovule is divided into two clusters (Supplementary Fig. 9, Supplementary Table 11). Interestingly, when we measured the Jaccard distance among all clusters (including the species with one female sample), we observed no grouping of similar clusters, indicating that the female gamete transcriptomes were poorly conserved (Supplementary Fig. 9). Functional enrichment analysis showed enrichment mainly for not assigned functions and RNA processing, and depletion for many biological processes (Supplementary Fig. 9). The *G. biloba* ovule cluster (GINBI-0, ovule) showed enrichment for many functions, but ovule samples of other species did not support this observation. Despite the small number of samples included these results provide evidence that female gamete transcriptomes are poorly conserved across the different species analyzed.

### Analysis of signaling networks underpinning male gametophyte development and function

Gene co-expression networks help to identify sets of genes involved in related biological processes and highlight regulatory relationships ^56,57^. Since we identified different gene clusters for male sub-samples (see above), we decided to test whether the genes assigned to different clusters are co-expressed. For this purpose, we reconstructed the co-expression networks of the ten species and analyzed whether the number of observed connections was similar to the number of expected connections (see material and methods). Interestingly, the clusters with expression profiles related to sperm had the least number of connections with other clusters for *O. sativa, Z. mays, A. trichopoda,* and *A. thaliana* (Fig. 6a). However, this pattern was not clear in *S. lycopersicum*, where the sperm cluster had connections with the cluster of generative cells. Specifically, for *A. thaliana* the co-expression network revealed that cluster C5 (sperm) is not well connected with other clusters (Fig. 6b), suggesting that the sperm cell transcriptome is distinctive, confirming earlier observations ^58–61^. The connections between clusters followed a pattern from cluster C0 to C4, which highlighted the interaction of genes among the different developmental stages of male gametogenesis. The number of transcription factors and kinases present in the co-expression network changed among the different clusters, where transcription factors seemed to be more abundant in cluster C0 (microspore), while kinases were more abundant in cluster C3 (mature pollen) (Fig. 6b, indicated by the sizes of rectangles, Supplementary Table 12).

**Fig. 6:**
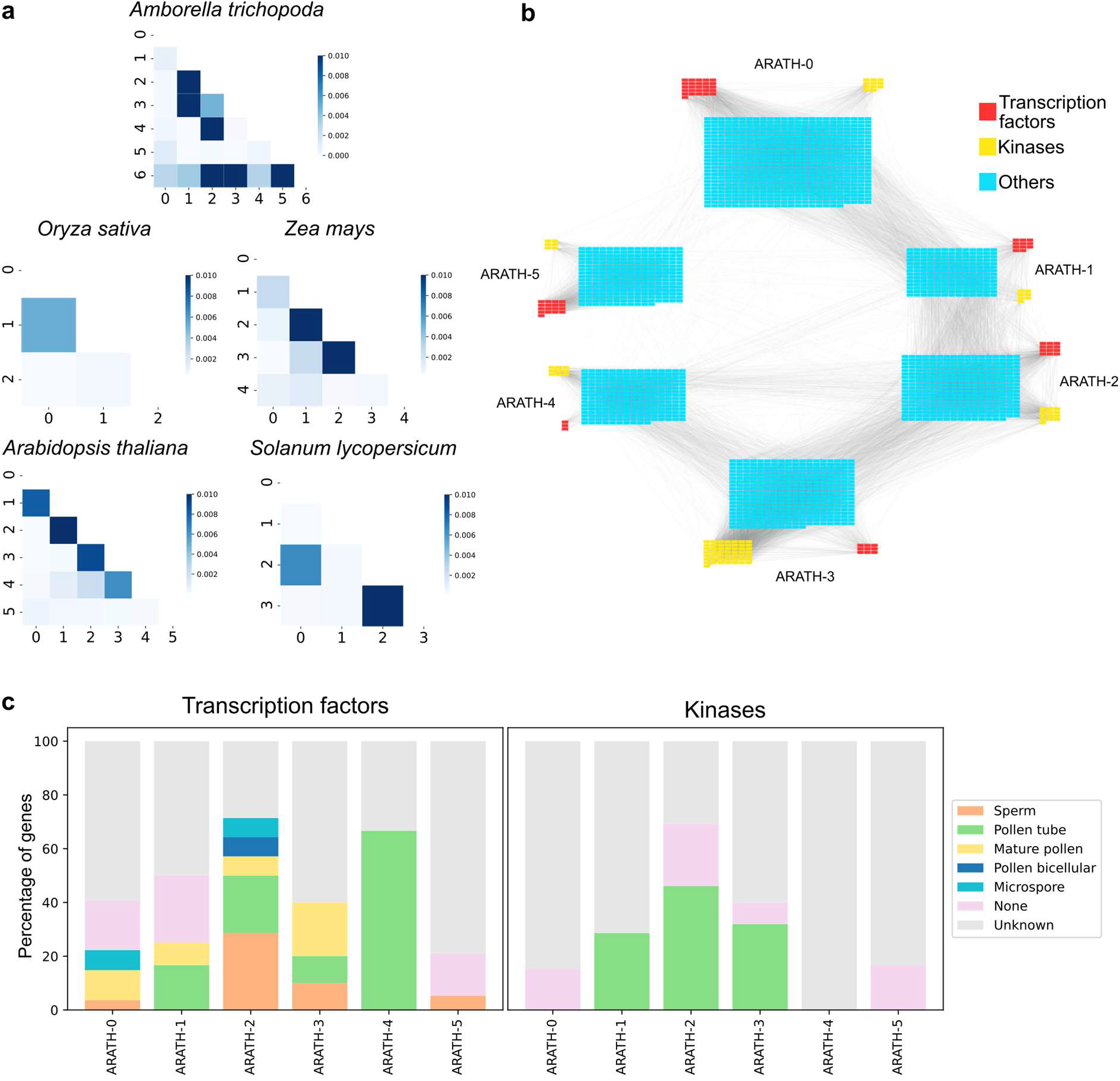
A network analysis of male clusters. **a**, Heatmaps show the number of observed connections divided by the number of expected connections. Darker colors indicate more connections between clusters. **b**, *A. thaliana* co-expression network clusters showing the edges between the different clusters (indicated as ARATH-0-5). The size of the panels indicate the number of genes in each cluster. Transcription factors, kinases, and other genes are shown in red, yellow, and blue, respectively. **c**, Percentage of genes of each *A. thaliana* male cluster. The colors indicate the different stages of male development that a given gene is known to be involved in. For example, the majority of transcription factors in cluster ARATH-4 (highest expression in the pollen tube, Fig. 5f) are important for pollen tube growth (green bars).

Transcription factors and kinases are regulatory proteins essential for plant growth and development. To uncover the regulatory mechanism underlying male gametogenesis, we analyzed all the predicted transcription factors and kinases in all the male clusters of *A. thaliana*. First, we searched for all the transcription factors and kinases present in the five clusters that have been characterized using experimental studies with mutants (Supplementary Table 13). Then we classified the effect of each mutant gene as follows: no effect related to male gametogenesis (none), no experimentally described function (unknown), and important for microspore, bicellular, mature pollen, pollen tube, and sperm function. Interestingly, most of the genes are described as unknown (Fig. 6c), indicating no experiments associated with those genes. It is important to note that the genes classified as ‘none’ have been found to have an effect in other organs, but since pollen phenotype can be easily missed, this does not rule out the possibility of these genes being associated with male development. Also, many of those genes show effects in roots, and it has been shown that some genes are active during tip growth of root hairs and pollen tubes ^62^. We observed that the transcription factors were important at different stages of male development, with main phenotypes affecting pollen tube and sperm function. Conversely, kinases only showed an effect on pollen tubes, which is in line with their intercellular communication involvement. Interestingly, we observed that genes present in the pollen tube cluster (ARATH-4) only affected pollen tube function, but pollen tube function can also be affected by genes from earlier stages of pollen development (ARATH1-3). In the case of sperm function, transcription factors expressed in tricellular pollen have the greatest effect, but we also observed the involvement of genes expressed in microspore, mature pollen and sperm (Fig. 6c).

### Comparative gene expression analyses with the EVOREPRO database

To provide easy access to the data and analyses generated by our consortium, we have constructed an online database available at www.evorepro.plant.tools. The database is preloaded with the expression data used in this study and also includes *Vitis vinifera* (eudicot, grapevine), *Chlamydomonas reinhardtii* (chlorophyte), and *Cyanophora paradoxa* (glaucophyte), bringing the total number of species to 13. The database can be queried with gene identifiers and sequences but also allows sophisticated, comparative analyses.

To showcase a typical user scenario, we identified genes specifically expressed in male organs (defined as, e.g., >35% reads of a gene expressed in male organs for Arabidopsis, Supplemental Figure 1). This can be accomplished for one (https://evorepro.sbs.ntu.edu.sg/search/specific/profiles) or two (https://evorepro.sbs.ntu.edu.sg/specificity_comparison/) species, where the latter option can reveal specific expression profiles that are conserved across species (Fig. 7a). For this example, we selected Arabidopsis and Amborella as species A and B from the drop-down menus, respectively, and used gene families comprising only land plants, which uses all species found under node 8 in the species tree (Fig. 3a). Alternatively, the user can also select gene families constructed with seed plants (11 species found under node 13, Fig. 3a) or archaeplastida (23 species found under node 1, Fig. 3a) sequences. Next, to select male organs for comparisons, we specified ‘Tissue specificity’ and ‘Male’ as a method to group the RNA-seq samples according to the definitions in Table 1. The slider near ‘SPM cutoff’ allows the user to adjust the SPM value (the slider ranges from SPM 0.5 to 1), which interactively reveals many genes are deemed organ-specific at a given SPM value cutoff. We left the slider at the default value (0.85) and clicked on the ‘Compare specificities’ button. The analysis revealed that 319 gene families are expressed specifically in the male organs of both Amborella and Arabidopsis (Fig. 7b), while the table below showed the identity of the genes and gene families (Fig. 7c, Table S15). Interestingly, among the conserved genes, we observed *GCS1/HAP2,* which is required for pollen tube guidance and fertilization ^63^. The table also contains links that redirect the user to pages dedicated to the genes and gene families. For example, clicking on the Arabidopsis *GCS1/HAP2* gene identifier redirects the user to a gene page containing the DNA/protein sequences (https://evorepro.sbs.ntu.edu.sg/sequence/view/17946), expression profile (Fig. 7d), gene family, co-expression network, and Gene Ontology functional enrichment analysis of the gene ^64^. As expected, the interactive, exportable expression profiles confirmed that the Arabidopsis *GCS1/HAP2* and the Amborella ortholog (https://evorepro.sbs.ntu.edu.sg/sequence/view/45084, Fig. 7e) are male-specific, with the highest expression in sperm and pollen. Clicking on the gene family identifier (OG_05_0008081) redirects to the gene family page (https://evorepro.sbs.ntu.edu.sg/family/view/139708), which among others, contains an interactive phylogenetic tree (Fig. 7f, https://evorepro.sbs.ntu.edu.sg/tree/view/88288) and heatmap (Fig 7g, https://evorepro.sbs.ntu.edu.sg/heatmap/comparative/tree/88288/row) showcasing the male-enriched expression profiles for most of the genes in this family. Therefore, this approach can be used to identify conserved, organ-specific genes across two species and study family-wide expression patterns.

**Fig. 7:**
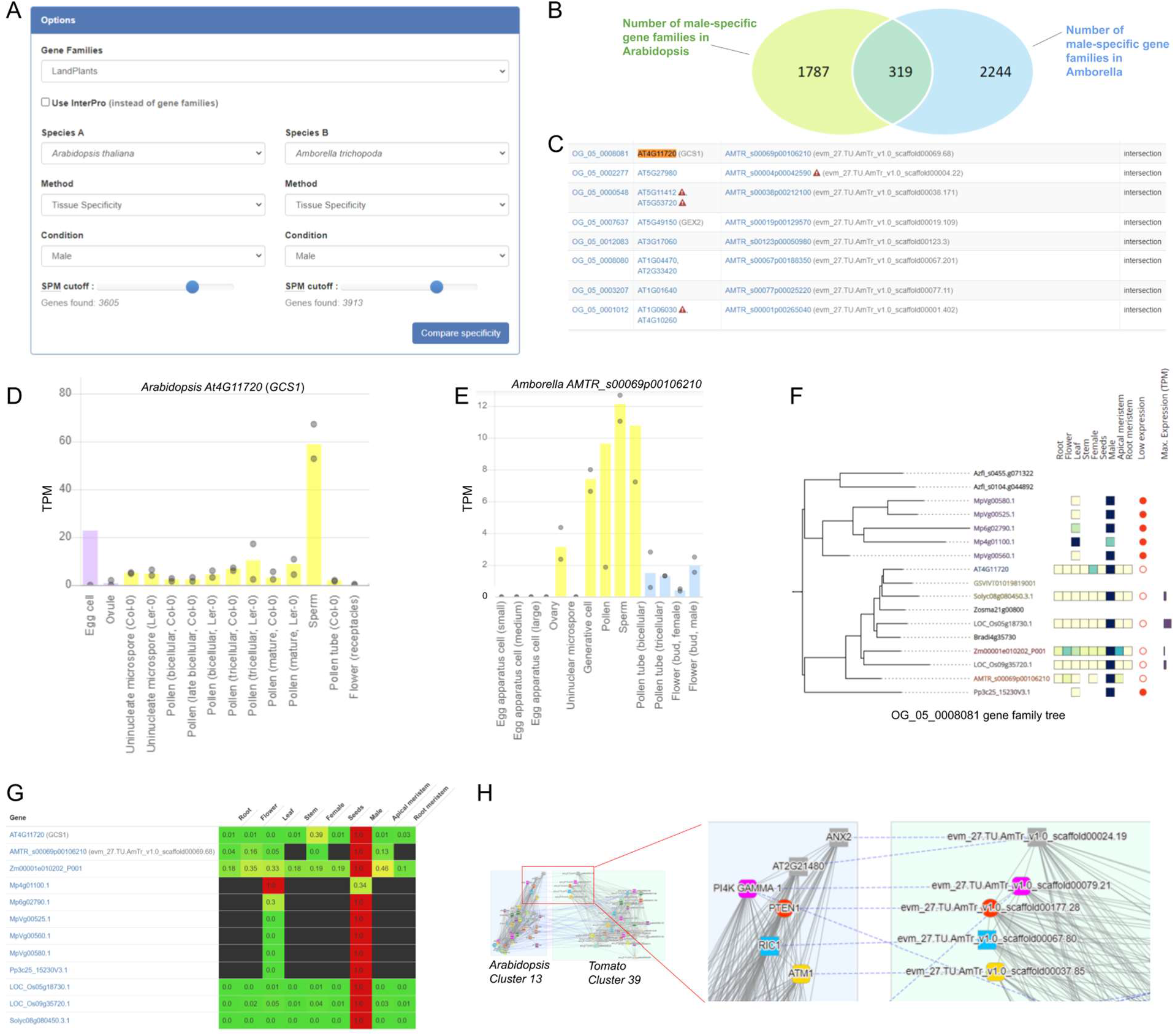
Features of the EVOREPRO database. **a**, Compare specificities tool. The dropdown menus allow selection of the species, gene families, organs, tissues, cell types, and SPM value cutoffs. The analysis is started by clicking on the ‘Compare specificity’ button. **b.** The Venn diagram shows the number of unique and common gene families of male-specific genes in Arabidopsis and Amborella. The default SPM value cutoff of 0.85 was used for both species. **c.** The table shows the identity of genes and gene families (first column) that are specifically expressed in male organs of Arabidopsis (second column) and Amborella (third column). Each row contains a gene family, and each cell can contain multiple comma-separated genes. Red triangles containing exclamation marks indicate genes with low expression (<10TPM). **d.** Expression profile of *GCS1* from Arabidopsis. The colored columns indicate the average expression values in the different samples, while gray points indicate the minimum and maximum expression values. The y-axis indicates the TPM value. **e.** Expression profile of *GCS1* -like gene from Amborella *(AMTR_s00069p00106210).* For clarity, the gray point indicating the maximum value in the sperm sample is omitted. **f.** Phylogenetic tree of the gene family OG_05_0008081 representing *GCS1.* The branches represent genes that are color-coded by species. The heatmap to the right of the gene identifiers indicates the scaled expression values in the major organ and cell types and ranges from low (yellow) to high (dark blue). Genes with TPM < 10 are indicated by filled red points, while the maximum gene expression is indicated by a blue bar to the right. **g.** Heatmap indicating the low (green) and high (red) expression of the *GCS1* gene family. **h.** Comparative analysis of co-expression clusters significantly (P<0.05) enriched for ‘pollen tube’ gene ontology term in Arabidopsis (cluster 13, left) and Amborella (cluster 39, right). Nodes indicate genes, while solid gray and dashed blue edges connect co-expressed and orthologous genes, respectively. We used ‘label co-occurrences’ as node options. For clarity, only part of each cluster is shown.

Alternatively, the database can be used to identify conserved co-expression clusters of functionally enriched genes. To demonstrate this tool, we navigated to https://evorepro.sbs.ntu.edu.sg/search/enriched/clusters and entered ‘pollen’ into GO text box, selected ‘pollen tube’ as query and clicked on ‘Show clusters’. The analysis revealed 5 co-expressed clusters significantly (P<0.05) enriched for ‘pollen tube’ gene ontology term in Arabidopsis. We clicked on one of the clusters (cluster 13, https://evorepro.sbs.ntu.edu.sg/cluster/view/113), redirecting us to a page dedicated to the cluster. As expected, the cluster is significantly (P<0.05) enriched for genes involved in pollen tube growth, cell wall organization and kinase activity, which are processes required to expand and direct the pollen tube to the ovule. The page contains the identity of the 152 genes found in this cluster, their average expression profiles, co-expression network (https://evorepro.sbs.ntu.edu.sg/cluster/graph/113), and gene families and protein domains found in the cluster.

Furthermore, a table labeled ‘Similar Clusters’ reveals the identity of similar (defined by Jaccard index, see methods) co-expression clusters in other species, which can be used to identify functionally equivalent clusters across species rapidly. To exemplify this, we first clicked on ‘Jaccard index’ table header to sort the similar clusters and clicked on the ‘Compare’ link next to Cluster 39 from Amborella (https://evorepro.sbs.ntu.edu.sg/graph_comparison/cluster/113/769/1). This redirected us to a co-expression network page showing the genes (nodes), co-expression relationships (gray edges), and orthologous genes (colored shapes of nodes connected by dashed edges) conserved in the two clusters. The analysis revealed many conserved genes essential for pollen function, such as *ANX2*^65^, *BUPS2 (At2g21480) ^66^, PI4K Gamma-1*^67^, *PTEN1 ^68^, RIC1*^69^, and *ATM1* ^70^. To conclude, this approach can be used to uncover functionally equivalent, conserved transcriptional programs.

## Discussion

To study the evolution of plant organs and gametes, we have generated and analyzed gene expression for ten land plants, comprising representatives of bryophytes, lycophytes, gymnosperms, basal angiosperms, monocots and eudicots. Our analyses’ main advantage is that the conclusions are drawn from comparative analyses of ten species, which cover the largest collection of representatives of land plants. The comparative analysis revealed that each organ type typically expressed >50% of genes, with the exception of the male gametes, which showed expression of ~38% of genes, on average (Figure 1D). Conversely, male gametes and roots showed the highest number (5.3% and 5.0%, respectively) of specifically expressed genes (Figure 1F), suggesting that these non-photosynthesizing cell types and tissues are highly unique and specialized.

With the surprising exception of female gametes, the corresponding transcriptomes tend to be more similar across the analyzed samples (Figure 2D, Figure S3, Figure S4). Another exception is seen in the leaf-like organs of bryophytes (leaflets and thallus for *Physcomitrella* and *Marchantia,* respectively), indicating that these organs have evolved independently from the leaves of flowering plants or that they have significantly diverged since the last common ancestor of flowering plants and bryophytes.

Next, we examined expression patterns of expressed gene families as a function of their age. We report a clear trend of older gene families having more ubiquitous (i.e., less organ-specific) expression, while younger gene families show an increasingly higher proportion of organ-specific expression (Figure 3b-c). This indicates that newly-acquired genes are typically recruited to perform some specialized function in a plant organ, tissue, or cell type, rather than being integrated into fundamental biological pathways. As expected, male gametes show the highest expression of the youngest genes (Figure 3d-e, Figure S7), which is in line with previous studies ^42,71^. Interestingly, *Physcomitrella* gametes did not show this pattern, which is a finding that warrants further studies.

To study how new functions were gained or lost as the organs and gametes evolved, we studied which phylostrata are enriched or depleted in the different organs (Figure 4a). Interestingly, we observe a significant enrichment for gene families that appeared long before the corresponding organ (Figure 4a), showing that the establishment of organs relies heavily on the co-option of existing genetic material, as suggested previously ^45,50^. Flowers (appearance in angiosperms), stems (appearance in vascular plants) and roots (appearance in vascular/seed plants) show similar patterns of enrichment and depletion of genes (Fig. 4a). This is surprising, as these organs appeared at different stages of plant evolution, which suggests that the co-option underlying the establishment of novel organs follows a similar pattern of gene gains and losses. Based on the diverse patterns of gains and losses of organ-specific gene families (Figure 4b) we conclude that monocot-specific families show substantial net gains in genes that are specifically expressed in male gametes, seeds, stems, roots or in apical and root meristems (Figure 4b), suggesting that during monocots evolution organ-specific transcriptomes was enriched with novel functions. Surprisingly, eudicots show an opposite pattern, exhibiting more net losses of organ-specific families in flowers, female and male gametes, leaves, stems, roots, and apical meristems (Figure 4b). This surprising pattern of loss of functions in eudicots merits investigation by further analysis, which is made possible by identifying the corresponding gene families (Table S8) and genes (Table S6).

Our comparative analysis of male gamete development reveals that transcriptional programs of mature pollen form well-defined clusters and are thus conserved across species (Figure 5f-g). The mature pollen clusters are enriched for processes related to signaling (protein modification comprising protein kinases) and cell wall remodeling (Figure 5e), which are likely representing processes mediating pollen germination, pollen tube growth, and sperm cell delivery. Conversely, the earlier stages of male gamete development showed less defined clusters and enrichment for genes with unknown function (bin ‘not assigned’, Figure 5e), suggesting that the processes taking place in the early stages of pollen development are yet to be uncovered. Furthermore, the female gametes show poor clustering, indicating overall low conservation of the transcriptional programs and enrichment of genes with unknown function for most clusters (Figure S9c). These results indicate that genes expressed during early male gamete and female gamete formation warrant closer functional analysis, which is now made possible by our identification of these genes (Table S10-11).

Of particular interest are the male-specific transcription factors and kinases that we identified (Figure 6c), assumingly involved in various stages of pollen development and function (Table S13). As a large fraction of these genes are not yet characterized, their involvement in male gametogenesis and function should be further investigated.

To provide easy access to the 13 expression atlases, organ-specific genes, functional enrichment analyses, co-expression networks, and various comparative tools, we provide the EVOREPRO database (www.evorepro.plant.tools) to the community (Figure 7). This database represents a valuable resource for further study and validation of key genes involved in organogenesis and land plants reproduction.

## Methods

### Physcomitrella growth conditions, RNA isolation and sequencing

#### Plant growth

The Gransden wild-type strain from *P. patens* Bruch & Schimp ^72^ was used for this study. To initiate plant growth and culture, 3 mature sporophytes were sterilized using a 5% commercial bleach solution for 5 minutes and rinsed twice in molecular grade water. Sterilized sporophytes were then broken using a pipette tip and diluted into 5mL molecular grade water. Spore containing solution was then distributed into 4 sterile peat pellets (Jiffy-7, Jiffy Products International) and two 9 cm Petri dishes containing KNOPS medium (Reski and Abel, 1985) supplemented with 0.5 g/l ammonium tartrate dibasic (Sigma-Aldrich Co). Petri dishes were kept at 25°C, 50% humidity, and 16 h light (light intensity 80 μlum/m/s). Protonema samples were collected 10 days after spore germination.

Plants in Phytatray™ II (Sigma-Aldrich Co) containing 4 sterile peat pellets (Jiffy-7, Jiffy Products International) were grown for 6-8 weeks at 25°C, 50% humidity, and 16 h light (light intensity 80 μlum/m/s). Water was supplied to the bottom of each box. Leave samples were collected after 6 weeks, prior to induction of gametangia development. For gametangia and sporophyte development, water was again supplied to the bottom of each box containing four pellets and were transferred to 17°C, 8 h light, and 50% humidity (light intensity 50 μlum/m/s) to induce the development of reproductive structures ^73^. Gametangia samples (archegonia, paraphysis and sperm cell packages) were collected 15 days after reproductive induction. Antheridia samples were collected at several time points during their development. Further development of the sporophyte was conducted under these conditions and sporophyte samples were collected at different time points during sporophyte development. S1 sporophytes were collected 7 days after sperm cell (SC) release, S2 sporophytes 15 days after SC release, S3 sporophytes 20 days after SC release (green spore capsules) and SM samples 28 days after SC release (brown spore capsules).

#### Sample preparation and sequencing

Leaves, protonema and sporophytes were collected under a stereoscope using tweezers, placed in 2.5 uL of RLT+ buffer (Qiagen), and shock frozen in liquid nitrogen. Before RNA-seq library preparation, these samples were mechanically disrupted using sterile pellet pestles (Z359947, Sigma-Aldrich Co). Antheridia, archegonia, paraphysis and sperm cell packages were collected using a Yokogawa CSU-W Spinning Disk confocal with 10x 0.25NA objective, using the brightfield channel and an Andor Zyla 4.2 sCMOS camera. For each of these samples the plants were prepared under a stereoscope, isolating the whole gametangia for ca. 10 shoots. They were placed in 20 uL of molecular grade water on a glass slide. Using a cover slip the gametangia were disrupted into individual antheridia by applying slight pressure. Slides were placed under a microscope and specific organs were identified and collected, using an Eppendorf CellTram® Air/Oil/vario micromanipulator with glass capillaries (borosilicate glass with fire polished ends, without filament GB100-9P) pulled with a Narishige PC-10 puller. Then they were transferred to another clean slide, and subsequently excessive liquid containing possible contaminations, such as cell debris, was removed. For paraphysis samples 8-15 individual paraphysis were collected directly into 2 uL of RLT+ buffer and flash frozen in liquid nitrogen. For antheridia samples 5 to 20 individual antheridia of each specific stage (9 to 15 days after induction, distinguished by size) were collected and then burst under a microscope by applying pressure on a cover slip applied to the samples on the slide. The slide was washed with 4 uL of RLT+ buffer and the buffer transferred into a PCR tube, subsequently flash frozen. Archegonia samples were prepared from 3-5 archegonia following the same procedure. Released sperm cell packages (2-5 per sample) were collected from gametangia preparations (as described above; antheridia 15 days after induction) without clean up, transferred into a tube with 2 uL of RLT+ buffer, flash frozen in liquid nitrogen and subsequently used for RNA-seq library preparation.

RNA-seq library preparation for all samples was performed as described in ^74^, with the addition of mixing the PCR tubes on a Thermomixer C (Eppendorf) every 15 minutes at 200 rpm for 1 min during the RT step. Libraries were sequenced on a NextSeq500 instrument with single-end 75 bp read length (SE75).

#### Marchantia growth conditions, RNA isolation and sequencing

Male accession of *Marchantia polymorpha* L., Takaragaike (Tak)-1 was grown on vermiculite under a long-day condition (16/8 h day/night) at 22 °C. To induce sexual reproduction, thalli developed from gemmae were transferred to a far-red light (700 – 780 nm, 44.3 μmol photons m^-2^s^-1^) supplemented light condition using LabLEDs (RHENAC GreenTec Ag). Sperms were released from antheridiophores by applying ddH2O supplemented with RNasin® Ribonuclease Inhibitor (1 u/μl, Promega), collected in a 1.5 mL tube, and pelleted by centrifugation at 3,000 *g* for 5 min at 4 °C. RNA-seq libraries were generated from total RNA of isolated *M. polymorpha* sperm using Smart-seq2 ^75^ using independent biological replicates. The libraries were sequenced on an Illumina Hiseq 2500 using 125 bp paired-end.

### Amborella growth conditions, RNA isolation and sequencing

#### Plant material and isolation procedures

*Amborella trichopoda* male flowers were harvested from a male plant growing in the Botanical Garden in Bonn (Germany), in a shaded place inside a greenhouse under controlled conditions of 16-18°C, constant humidity of 66% and 12-hour photoperiods. Buds and fully opened male flowers were gathered in 50 ml Falcon™ conical tubes (Thermo Fisher), placed without lid in a hermetically sealed plastic box containing a bed of silica gel.

Uninucleated microspores (UNM) were isolated at room temperature from flower buds of 4.5 mm length, as these were found to contain 98% uninucleated microspores. In brief, three samples with each 5 g buds were homogenized in 0.1 M mannitol and filtered with a 70-micron pore size PET strainer (PluriSelect). The filtered solution was processed by subsequent steps of percoll gradient separation, washing and centrifugation, as described previously ^76^.

*Amborella* generative cells (GC) were obtained from mature pollen grains that were purified like described previously ^77^. Per replicate, 50 mg pollen was resuspended in 1 ml pollen germination medium and transferred into a 1.5 ml vial containing glass beads (0.4 – 0.6 mm). The vial was vortexed continuously at 2,200 rpm for 4 minutes to crack the pollen grains and release its contents. The solution was filtered using a 15-micron PET strainer (PluriSelect). To stain the nuclei, a final concentration of 10X SYBR Green I was added and GCs were identified using an inverted microscope (Nikon) equipped with high-resolution 20X and 40X objectives suitable for fluorescent applications and suitable filters for SYBR Green I (497 nm excitation; 520 nm emission). For RNA-seq, three replicates of each 140 GC were harvested manually using an Eppendorf CellTram.

*Amborella* sperm cells (SC) were isolated at room temperature by adapting a method described for tomato sperm cell isolation ^78^. In brief, three replicates with each 50 mg purified pollen were germinated as described ^77^. 16 hours after germination, the medium was removed by filtration using a 15-micron PET strainer (PluriSelect) and the pollen tubes were incubated for 10 min in a 15% mannitol solution with 0.4% cellulase “Onozuka” R-10 and 0.2% macerozyme R-10 to release the sperm cells. The mixture was refiltered using a 15-micron PET strainer and loaded on 5 ml 23% Percoll in 0.55 M mannitol and centrifuged at 1,000 x g for 30 min. Approximately 1 ml with SC, floating on the surface of the Percoll gradient, were harvested, washed with 1 ml RNAprotect® Cell Reagent (Qiagen) and centrifuged for 10 min at 2,500 x g. 50 μl of SC-enriched pellet (approximately 250 sperm cells each replicate) was used for RNA-seq library preparation.

Isolation and sampling of *Amborella* ovaries, egg apparatus cells, pollen tubes, pollen grains as well as male and female flowers, tepals, roots and leaves was done as described in previous studies ^77,79^.

#### RNA isolation and sequencing

RNA isolation from uninucleated microspores was performed by using the Spectrum™ Plant Total RNA Kit (Sigma-Aldrich) according to manufacturer’s instructions. Total RNA from *Amborella* generative cells and sperm cells was extracted according to the “Purification of total RNA from animal and human cells” protocol of the RNeasy Plus Micro Kit (QIAGEN, Hilden, Germany). In brief, cells were stored and shipped on dry ice. After adding RLT Plus containing ß-mercaptoethanol the samples were homogenized by vortexing for 30 sec. Genomic DNA contamination was removed using gDNA Eliminator spin columns. Next ethanol was added and the samples were applied to RNeasy MinElute spin columns followed by several wash steps. Finally total RNA was eluted in 12 μl of nuclease free water. Purity and integrity of the RNA was assessed on the Agilent 2100 Bioanalyzer with the RNA 6000 Pico LabChip reagent set (Agilent, Palo Alto, CA, USA).

The SMARTer Ultra Low Input RNA Kit for Sequencing v4 (Takara) was used to generate first strand cDNA from 2.5 ng UNM, 0.8 ng GC and 0.5 ng SC total RNA. Double stranded cDNA was amplified by LD PCR (10 for UNM, 13 cycles for GC and 15 cycles for SC) and purified via magnetic bead clean-up. Library preparation was carried out as described in the Illumina Nextera XT Sample Preparation Guide (Illumina, Inc., San Diego, CA, USA). 150 pg of input cDNA were tagmented by the Nextera XT transposome. The products were purified and amplified via a limited-cycle PCR program to generate multiplexed sequencing libraries. For the PCR step 1:5 dilutions of index 1 (i7) and index 2 (i5) primers were used. The libraries were quantified using the KAPA SYBR FAST ABI Prism Library Quantification Kit. Equimolar amounts of each library were used for cluster generation on the cBot (TruSeq SR Cluster Kit v3). The sequencing run was performed on a HiSeq 1000 instrument using the indexed, 2=100 cycles paired end (PE) protocol and the TruSeq SBS v3 Kit. Image analysis and base calling resulted in .bcl files, which were converted into .fastq files by the CASAVA1.8.2 software. Library preparation and RNA-seq were performed at the service facility “Center of Excellence for Fluorescent Bioanalytics (KFB)” (Regensburg, Germany; www.kfb-regensburg.de).

### Arabidopsis growth conditions, RNA isolation and sequencing

*Arabidopsis thaliana* accession Columbia-0 (Col-0) plants were grown in controlled-environment cabinets at 22°C under illumination of 150 μmol/m2/sec with a 16-h photoperiod. Mature pollen grains (MPG) were harvested from open flowers of 5 to 6-week old plants by shaking into liquid medium (0.1 M D-mannitol) as described previously^79^. Microspores and developing pollen grains were released from anthers of closed flower buds and purified by Percoll density gradient centrifugation as described ^76,80^. Populations of spores at five stages of development were isolated: uninucleate microspores (UNM), bicellular pollen (BCP), late bicellular pollen (LBC), tricellular pollen (TCP) and mature pollen (MPG).

For semi in vivo pollen tube growth, a transgenic marker line harboring MGH3p::MGH3-eGFP and ACT11p::H2B-mRFP ^21^ was used to pollinate WT emasculated pistils. After 2 hours, the pollinated pistil was excised and placed on double sided tape. The excised pistil was then cut at the junction of style and ovary and placed gently on solidified agarose pollen germination medium ^81^. The pistil was incubated for an additional 4 hours for the pollen tubes to emerge from the cut end of the style. The pollen tubes were harvested using a 25G needle and immediately frozen in liquid nitrogen and subsequently used for the RNA-seq library preparation as described in ^74^.

Total RNA was isolated from each sample using the RNeasy Plant Kit (Qiagen) according to the manufacturer’s instructions. RNA was DNAse-treated (DNA-freeTM Kit Ambion, Life Technologies) according to the manufacturer’s protocol. RNA yield and purity were determined spectrophotometrically and using an Agilent 2100 Bioanalyzer. cDNA was prepared using a slightly modified SmartSeq2 protocol in which cDNA is synthesized from poly(A)+ RNA with an oligo(dT)-tailed primer ^75,82^. The final libraries were prepared using a low-input Nextera protocol ^83^. Libraries were sequenced on a NextSeq500 instrument with single-end 75 bp read length (SE75).

A transgenic line expressing EC1.1p:NLS-3xGFP was cultured and used for Arabidopsis egg cell isolation as previously described ^84^. Three replicates of 25 to 30 pooled egg cells were used for RNA extraction, RNA-seq library preparation and Illumina Next Generation Sequencing ^85^.

### Tomato growth conditions, RNA isolation and sequencing

*Solanum lycopersicum* (tomato accession Nagcarlang, LA2661) seeds were obtained from the Tomato Genetics Resource Center (TGRC, https://tgrc.ucdavis.edu/) and grown in the Brown University Greenhouse (Providence, RI, USA). Dry pollen grains were collected from stage 15 flowers ^86^ into 500μl eppendorf tubes. Pollen tubes were grown in 300μl of pollen growth medium in a 750μl eppendorf tube that was incubated in a 28°C water bath. Pollen tubes were grown at a density of ~1000 pollen grains/μl. The pollen germination medium ^87^ comprised 24% (w/v) polyethylene glycol (PEG) 4000, 0.01% (w/v) boric acid, 2% (w/v) Suc, 20 mm MES buffer, pH 6.0, 3 mm Ca(NO3)2·4H2O, 0.02% (w/v) MgSO4·7H2O, and 1 mm KNO3. Pollen tubes were grown for 1.5 hours, 3 hours, or 9 hours before they were collected by centrifugation (1000 x g) for 1 minute. Pollen germination medium was carefully removed by pipetting to avoid disrupting the loose pollen tube pellet. Independent pollen collections were made for each of three biological replicates at each time point. Eppendorf tubes containing pollen tubes were immediately flash frozen in liquid N2, then stored at −80°C, or put directly on a dry-ice cooled metal block for cell disruption by grinding with a frozen plastic pestle (Kontes). Total RNA was extracted using the RNeasy Plant Kit (Qiagen). RNA samples were evaluated by Agilent 2100 Bioanalyzer (Brown University Genomics Core Facility) before RNA-seq library preparation (polyA selection) and Illumina HiSeq, (150bp, paired end) sequencing were performed by Genewiz (South Plainfield, New Jersey. USA).

### Maize growth conditions, RNA isolation and sequencing

Maize plants (inbred line B73) were grown in an air-conditioned greenhouse at 26°C under illumination of about 400 μmol/m2/sec with a 16-h photoperiod (21°C night temperature) and air humidity between 60-65%. Fresh mature pollen grains were harvested as described^88^. Pollen tubes were germinated and grown for 2 hours *in vitro* using liquid pollen germination medium ^89^. Total RNA was extracted from each three biological replicates of 100 mg pollen grains/pollen tubes by using a Spectrum™ Plant Total RNA Kit (Sigma-Aldrich) according to manufacturer’s instructions. 250 ng of total RNA was each used for library construction. RNA-seq was carried out as described in the Illumina TruSeq Stranded mRNA Sample Preparation Guide for the Illumina HiSeq 1000 System (Illumina) and the KAPA Library Quantification Kit (Kapa Biosystems). Data from sperm cells, egg cells and various zygote stages were taken from published data ^88^.

### Compiling gene expression atlases

RNA data of different samples from nine species (*Physcomitrium patens, Marchantia polymorpha, Ginkgo biloba, Picea abies, Amborella trichopoda, Oryza sativa, Zea mays, Arabidopsis thaliana, Solanum lycopersicum*) were grouped in ten different classes (flower, female, male, seeds, spore, leaf, stem, apical meristem, root meristem, root) (Table 1, Supplementary Table 1). For male and female reproductive organs samples we also included different sub-samples (female: egg cell, ovary, ovule; Male: microspore, bicellular pollen, tricellular pollen, mature pollen, pollen tube, generative cell, sperm) for each species (Table 1, Supplementary Table 1). A total of 4,806 different RNA sequencing samples were used, from which 4,672 were downloaded from the SRA database and 134 obtained from our experiments (see above). Publicly available RNA-seq experiments data were downloaded from ENA ^90^, as described in CoNekt-Plants ^64^. Proteomes and CDSs of each species were downloaded from different sources (Supplementary Table 14). The raw reads of each sample were mapped to the coding sequences (CDS) with Kallisto v.0.46.1 ^26^ to obtain transcripts per million (TPM) gene expression values. If the reads came from single cell samples (egg cell, ovule, sperm, generative cell), we removed the samples that have <1M reads mapped, and for the other samples we removed those with <5M reads mapped (Supplementary Table 1). All those samples were used to calculate Highest Reciprocal Rank (HRR) networks, where two genes with HRR<100 were connected ^91^. For comparative expression analysis, an additional filter was applied by keeping only samples with a Pearson correlation coefficient (PCC) >=0.8 to at least one other sample of the same type (e.g. flower to flower) (Supplementary Table 1). Additionally, we included the expression matrix of *Selaginella moellendorffii* which has 18 samples (Supplementary Table 1), and exclusively for the database (see section Constructing the co-expression network and establishing the EVOREPRO database) the expression matrices of two unicellular algae *(Chlamydomonas reinhardtii* and *Cyanophora paradoxa)* and *Vitis vinifera*^92^. Finally, genes with median expression levels >2 TPM were considered as expressed ^93^. All expression matrices are available for download from http://www.gene2function.de/download.html.

### Identifying sample-specific genes

Sample-specific genes based on expression data were detected by calculating the specificity measure (SPM), using a similar method as described in ^94^. For each gene, we calculated the average TPM value in each sample (e.g., root, leaf, seeds). Then, the SPM value of a gene in a sample was computed by dividing the average TPM in the sample by the sum of the average TPM values of all samples. The SPM value ranges from 0 (a gene is not expressed in a sample) to 1 (a gene is fully sample-specific). To identify sample-specific genes, for each of the ten species, we first identified a SPM value threshold above which the top 5% SMP values were found (Supplementary Fig. S1, red line). Then, if a gene’s SPM value in a sample was equal to or larger than the threshold, the gene was deemed to be specifically expressed in this sample.

### Similarity of sample-specific transcriptomes between samples and species

To estimate whether sample-specific transcriptomes (see above) are similar, we calculated Jaccard distance *d_j_* between orthogroup sets. These orthogroup sets were found by identifying the orthogroups of sample-specific genes per each species. Then pairwise *d_j_* was calculated for all the samples and used as input for the clustermap. The *d_j_* ranges between 0 (the two sets of orthogroups are identical) to 1 (the two sets have no orthogroups in common).

To estimate whether a species’ sample-specific transcriptome was significantly similar to a corresponding sample in the other species (e.g. Arabidopsis root vs. rice root, tomato root), we tested whether the *d_j_* values comparing the same sample were smaller (i.e. more similar) than *d_j_* values comparing the sample to the other samples (e.g., Arabidopsis root vs. rice flower, rice leaf, tomato flower, tomato leaf). We used Wilcoxon rank-sum to obtain the p-values, which were adjusted using a false discovery rate (FDR) correction ^95^.

### Phylogenomic and phylostratigraphic analysis

We used proteomes of 23 species representing key phylogenetic positions in the plant kingdom (see Supplementary Table 14), to construct orthologous gene groups (orthogroups) with Orthofinder v2.4.0 ^96^, where Diamond v0.9.24.125 ^97^ was used as sequence aligner. A species tree based on a recent phylogeny including more than 1000 species ^98^ was used for the phylostratigraphic analysis. The phylostratum (node) of an orthogroup was assessed by identifying the oldest clade found in the orthogroup ^99^ using ETE v3.0 ^100^. To test whether a specific phylostratum is enriched in a sample, we randomly selected (without replacement) the number of observed sample-specific genes 1000 times. The empirical p-values were obtained by calculating whether the observed number of gene families for each phylostratum was larger (when testing for enrichment) or smaller than (testing for depletion) than the number obtained from the 1000 sampling procedure. The p values were FDR corrected ^95^.

### Transcriptomic age index calculation

Transcriptome age index (TAI) is the weighted mean of phylogenetic ranks (phylostrata) and we calculated it for every sample ^71^. We used the species tree from ^98^. The nodes in the tree were assigned numbers ranging from 1 (oldest node) to 22 (youngest node, Fig. 3a) by traversing the tree using ETE v3.0 (Huerta-Cepas et al. 2016) with default parameters. The age (phylostratum) of an orthogroup and all genes belonging to the orthogroup, were derived by identifying the last common ancestor found in the orthogroup using ETE v3.0 ^100^. In the case of species-specific orthogroups the age of the orthogroup was assigned as 23. Finally, all genes with TPM values <2 were excluded and the TAI was calculated for the remaining genes by dividing the product of the gene’s TPM value and the node number by the sum of TPM values.

### Functional annotation of genes and identification of transcription factor and kinase families

The proteomes of the ten species included in the transcriptome dataset were annotated using the online tool Mercator4 v2.0 (https://www.plabipd.de/portal/web/guest/mercator4/-/wiki/Mercator4/recent_changes). This tool assigns Mapman4 bins to genes ^101^. Transcription factors and kinases were predicted using iTAK v1.7a ^102^. Additional transcription factors were identified using the online tool PlantTFDB v5.0 (http://planttfdb.cbi.pku.edu.cn/prediction.php)^103^.

### Functional enrichment analysis

Functional enrichment of the list of sample-specific and cluster-specific genes of each species, and genes gained in each node, was calculated using the bins predicted with Mercator 4 v2.0. Briefly, for a group of *m* genes (e.g., genes specifically expressed in Arabidopsis root), we first counted the number of Mapman bins present in the group, and then evaluated if these bins were significantly enriched or depleted by calculating an empirical p-value. The empirical p-value that tests whether a Mapman bin (term) is enriched in a collection of *m* genes is defined as:

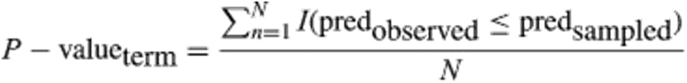

*Wherepred_observed_*is the number of times a term is observed, *pred_Sampled_* is the number of times the term is observed when the terms of *m* genes are randomly sampled (without replacement) from the all genes in the genome. *N* is the number of permutations, which was set to 1000. *I* is an indicator function, which takes a value of 1 when the event (in this case *pred_observed_* ≤ *pred_Sampled_*) is true, and 0 when it is not. For functional depletion analysis a similar approach was followed, with *I* taking a value of 1 when *pred_observed_ ≥ pred_sampled_*. To account for multiple hypothesis testing, we applied a false discovery rate (FDR) correction to the p-values ^95^. Transcription factor and kinase enrichment was calculated following the same procedure.

### Identification of orthogroup expression profiles

In order to analyse the expression profiles at phylostrata level, orthogroups were classified as ‘sample-specific’, ‘ubiquitous’, and ‘not conserved’. ‘Sample-specific’ orthogroups are orthogroups containing sample-specific genes and can be sub-classified according to the organ (flower-, female-, male-, seeds-, spore-, leaf-, apical meristem-, stems-, root meristem-, root-specific). ‘Ubiquitous’ are orthogroups that are expressed in different samples for each species, i.e., they do not show a ‘sample-specific’ expression profile. ‘Not conserved’ are orthogroups that have different sample-specific expression profiles in different species (e.g., orthogroups containing root-specific genes for *Arabidopsis* and male-specific genes for *Solanum*). Only orthogroups with species with sufficient expression data were used. More specifically, we only analyzed orthogroups that were: i) species-specific with transcriptome data or, ii) contained at least two species with transcriptome data. To identify sample-specific orthogroups, we required, iii) >50% of genes of the orthogroup should support the expression profile, iv) >=50% of the species with transcriptome data present in the node should support the expression profile.

### Gene enrichment analysis per phylostrata

In order to analyse gene enrichment of specific samples across the different phylostrata in the species tree (Fig. 3a), we used all the sample-specific genes of the ten species included. For each species and for each defined sample (ubiquitous, flower, female, male, seeds, spore, leaf, stem, apical meristem, root meristem, root) we counted the number of genes present in each node of the species tree, and then evaluated if the number of sample-specific genes were significantly enriched or depleted by calculating an empirical p-value as described for functional enrichment analysis. Then, we evaluated each sample and counted the number of species that show significant enrichment/depletion (p<0.05) in each node of the species tree. We obtained an normalized value per each node by calculating the difference of species showing enrichment and species showing depletion and dividing it by the total number of species that show enrichment/depletion. These results were used to plot a heatmap using the seaborn python package ^104^.

### Gene family comparisons

For each sample-specific (flower, female, male, seeds, spore, leaf, stem, apical meristem, root meristem, root) and ubiquitous expression profiles we mapped loss and gain of organ-specific gene families onto the species tree (Fig. 3a). All the orthogroups classified as sample-specific (see above) were analysed independently and gain and loss was computed using the approach described in ^105^ with ETE v3.0 ^100^. Briefly, a gene family gain was inferred at the last common ancestor of all the species included in the family and a loss when a species did not have orthologs in the particular gene family. Groups of monophyletic species that have lost the gene were counted as one loss. Then, we collapsed the values of the nodes of the species tree to fit the different clades included (Fig. 4b), and we calculated the difference between the total gains and the total losses to obtain an absolute value for each node. The values of each expression profile were normalized dividing the values by the maximum absolute value in a way that we got a range from -1 to 1 (negative values for losses and positive values for gains). Finally, per each expression profile (ubiquitous, flower, female, male, seeds, spore, leaf, stem, apical meristem, root meristem, root) a graphical representation of the different clades showing the nodes with a intensity of color proportional to the normalized values of gains and losses was plotted using ETE v3.0 ^100^.

### Identification of gamete-specific transcriptional profiles by clustering analysis

We analyzed the male and female sample-specific genes and their different sub-samples (Supplementary Table 1), to identify transcriptional profiles by clustering analysis. For the clustering analysis we only included species with at least 2 subsamples (*Amborella trichopoda*, *Oryza sativa*, *Zea mays*, *Arabidopsis thaliana*, *Solanum lycopersicum*). The male samples were divided into: microspore, bicellular pollen, tricellular pollen, mature pollen, pollen tube, generative cell, and sperm cell for Angiosperms; and sperm for bryophytes. The female samples were divided into egg cell, ovary, and ovule. For each gene, the average TPM in each sub-sample was calculated, and the average TPM values were scaled by dividing with the highest average TPM value for the gene. The k-means clustering method from the sklearn.cluster package ^106^ was used to fit the scaled average TPM values to the number of clusters (k) ranging from 1 to 20. The optimal number of *k* for each species was estimated by using the elbow method, where *k* that produced a sum of squared distances<80% of k=1 was selected (Supplementary Fig. 10). Seaborn ^104^ python package was used for plotting the figures.

### Constructing the co-expression network and establishing the EVOREPRO database

Coexpression networks were calculated by using Highest Reciprocal Rank (HRR) value ^91^, which is a distance-based metric that ranges from 0 (two genes are strongly coexpressed) to 100 (two genes are weakly coexpressed). The networks were constructed by a CoNekT framework ^64^, which was also used to establish the EVOREPRO database available at www.evorepro.plant.tools. For each species, all the genes that were co-expressed in each male cluster were analysed to test whether the number of connections observed is similar to the expected number. For this, we divided the number of observed connections between the genes of two clusters (eg. cluster 1 and cluster 2) by the expected value (product of the number of genes in cluster 1 x number of genes in cluster 2). These values were used to perform a pearson correlation analysis and the results were presented in heatmaps. The networks present in the male clusters were visualized using Cytoscape v3.8.0 ^107^. The network files are available from www.evorepro.plant.tools/species/.

## Data availability

The fastq files are available for Arabidopsis (E-MTAB-9456), Amborella (E-MTAB-9190), Marchantia (E-MTAB-9457), Physcomitrella (E-MTAB-9466), maize (E-MTAB-9692) and tomato (E-MTAB-9725).

## Acknowledgments

I.J is supported by Singaporean Ministry of Education grant MOE2018-T2-2-053, while M.M is supported by NTU Start-Up Grant. ERA-CAPS EVO-REPRO I2163 to F.B.; ERA-CAPS-0001-2014 to J.D.B; ERA-CAPS EVO-REPRO DR 334/12-1 to S.S. and T.D. DH was supported by ERA-CAPS UK Biotechnology and Biological Research Council Grant BB/N005090 awarded to DT; M.B. was supported through the FWF Lise Meitner fellowship M1818. The Vienna BioCenter Core Facilities GmbH (VBCF) Plant Sciences Facility acknowledges funding from the Austrian Federal Ministry of Education, Science and Research and the City of Vienna. L.S was supported by CSF grant 17-23183S. C.M. and D.Ho. were supported by Czech Ministry of Education, Youth and Sport (LTC18034 and LTAIN19030) through the European Regional Development Fund-Project “Centre for Experimental Plant Biology”: No. CZ.02.1.01/0.0/0.0/16_019/0000738.The Genomics Unit of Instituto Gulbenkian de Ciência was partially supported by ONEIDA Project (LISBOA-01-0145-FEDER-016417) co-funded by FEEI - ”Fundos Europeus Estruturais e de Investimento” from “Programa Operacional Regional Lisboa 2020” and by national funds from FCT - ”Fundação para a Ciência e a Tecnologia”. C.S.M acknowledges a doctoral fellowship from FCT (PD/BD/114362/2016) under the Plants for Life PhD Program. J.D.B received salary support from FCT through an “Investigador FCT” position. MJ and JG were supported by a US National Science Foundation grant (IOS-1540019).

Help with sample generation: Lenka Záveská Drábková and David Reňák. Marchantia growth was performed by the Plant Sciences Facility at Vienna BioCenter Core Facilities GmbH (VBCF), member of the Vienna BioCenter (VBC), Austria. Maximilian Weigend, Cornelia Löhne and Bernhard Reinken (Botanical Garden of the University of Bonn, Germany) are acknowledged for providing *Amborella trichopoda* plant material. We acknowledge Devendra Shivhare for help with initial analysis of *Physcomitrium* expression data.

We would like to thank Debbie Maizels (http://www.scientificart.com) for the illustrations on Fig.1 and Fig. 5.

## Author Contributions

Conceived and designed the analysis: JDB, MM

Collected the data: ACL, MFT, SGP, CSM, IJ, LS, CM, DHo, DH

Contributed data or analysis tools: FB, MB, SS, TD, DT

Performed the analysis: IJ, CF, SP, ACL, MM

Wrote the paper: IJ, JDB, MM

## Supplementary information

**Supplementary Fig. 1:**
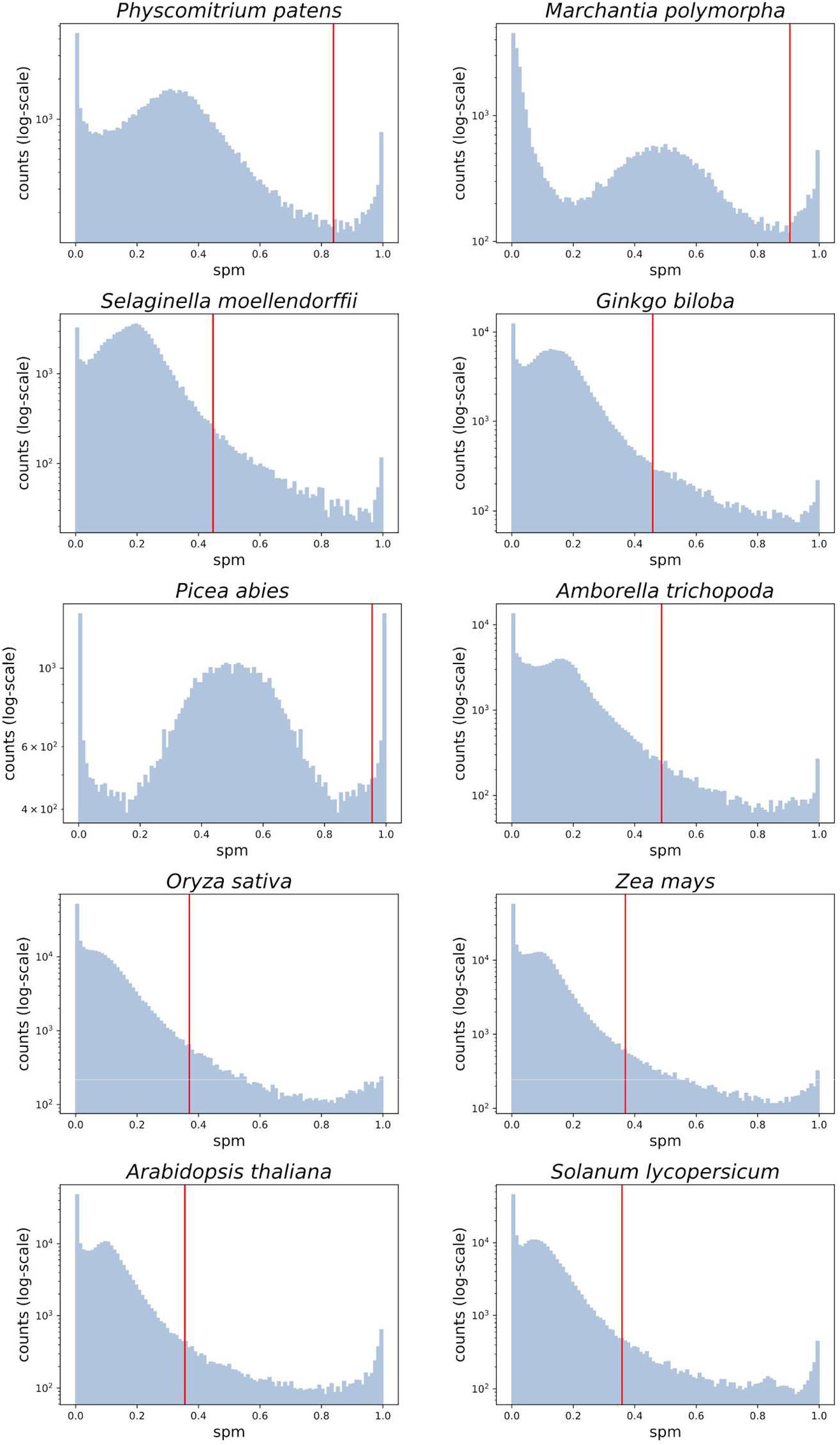
Distribution of SPM values in the ten species. The x-axis indicates the specificity measure (SPM), while the y-axis indicates the log10-transformed frequency of the SPM values observed for all genes across the samples. The vertical red line indicates the SPM value cutoff, below which 95% of values are found.

**Supplementary Fig. 2:**
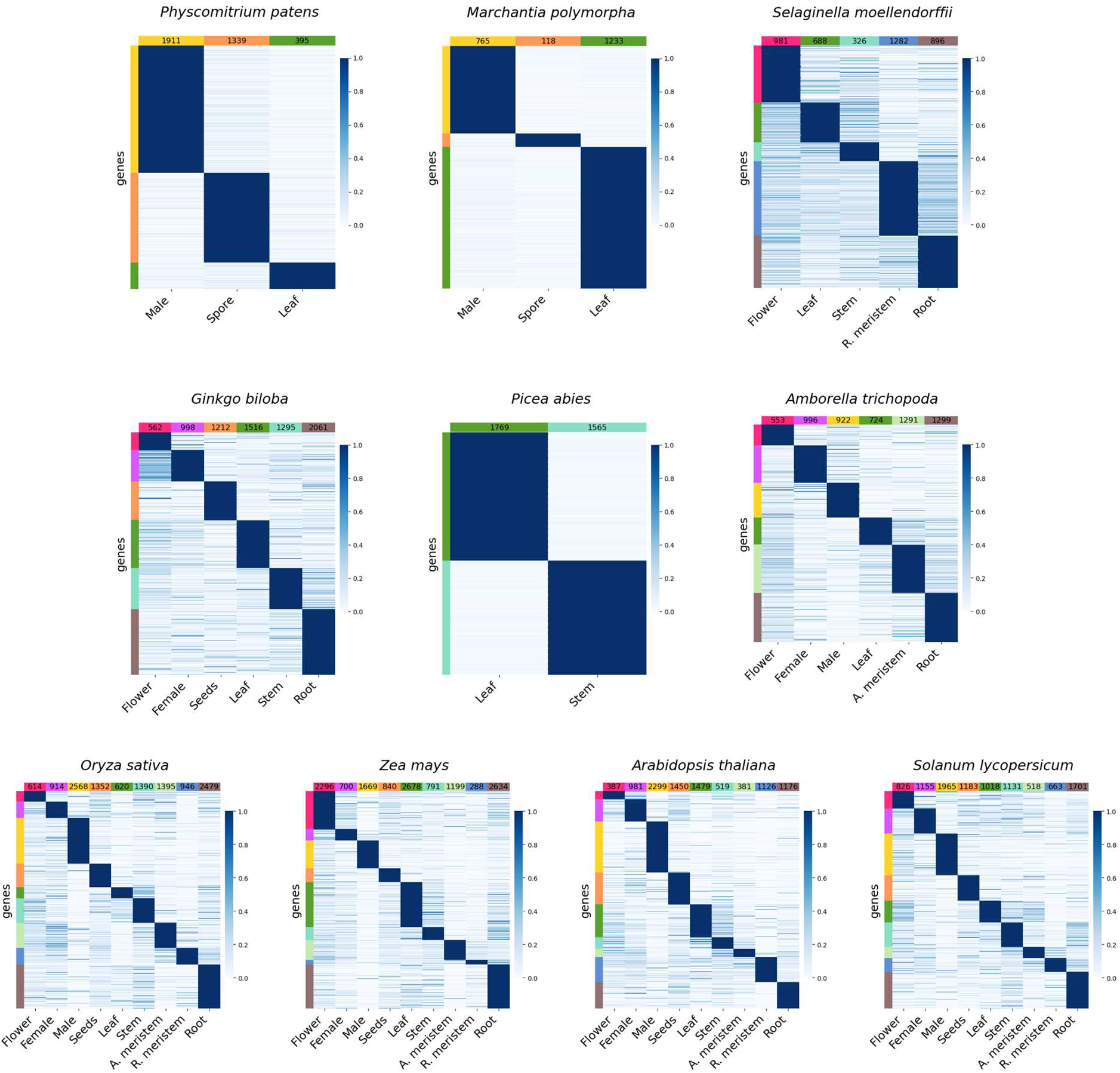
Expression profiles of the genes that were deemed to be specifically expressed in one of the organs/tissues/cells (sample) of the ten species used in this study. Genes are in rows, samples in columns, and the genes are sorted according to the expression profiles (e.g., flower, female). The numbers at the top of each column indicate the total number of specific genes in each sample. Gene expression is scaled to range from 0-1. Bars on the left of each heatmap show the sample-specific genes and correspond to the samples on the bottom: pink - Flower, purple - Female, yellow - Male, orange - Seeds/Spore, dark-green - Leaf, medium-green - Stem, light-green - Apical meristem, blue - Root meristem, brown - Root.

**Supplementary Fig. 3:**
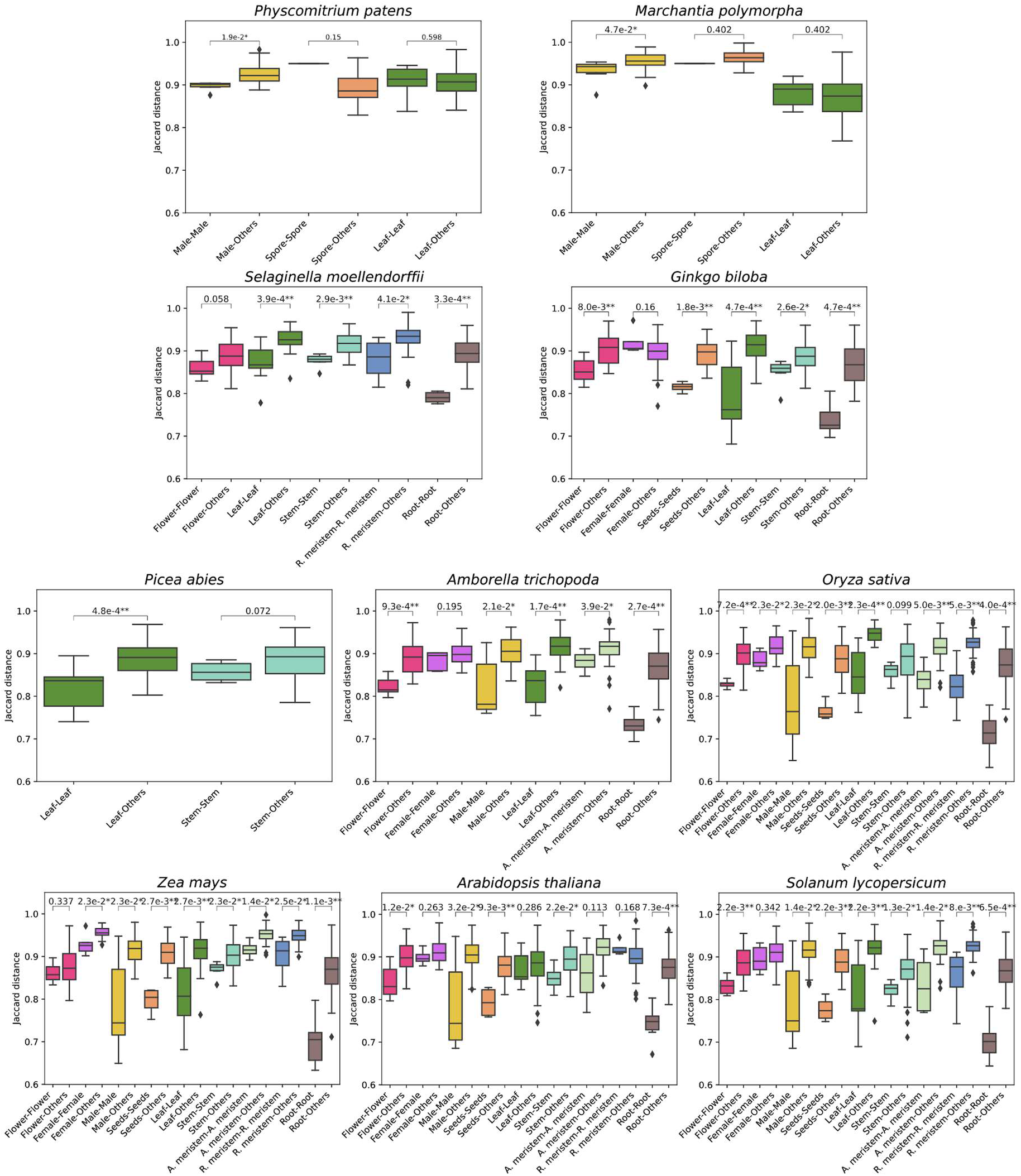
Bar plot showing the Jaccard distances when comparing the same samples (i.e., male-male) and one sample versus the others (i.e., male-others) for the ten species included in this study.

**Supplementary Fig. 4:**
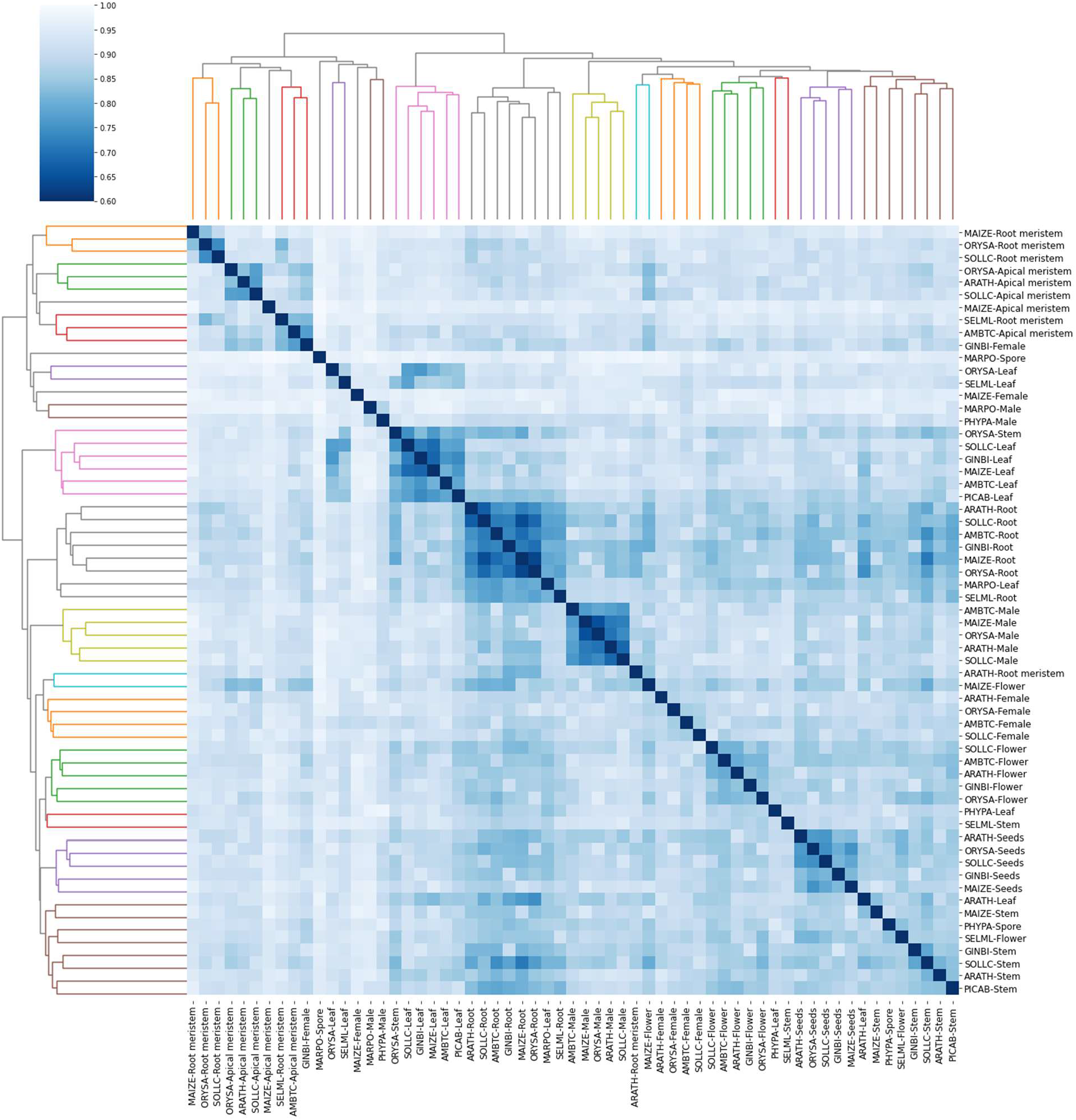
Comparing transcriptome similarities of the samples of the ten species. We used the Jaccard Index to calculate the similarity of transcriptomes of all samples in the dataset. The heatmap shows which transcriptomes of samples across species are similar by hierarchical clustering (dark blue). A lower value indicates a stronger similarity between two samples (white). For example, when comparing *Arabidopsis* root to roots from other species, we observe more similar transcriptomes than *Arabidopsis* root to non-root samples. The dendrograms on top and the left show the different clusters formed when the distance is <1.3.

**Supplementary Fig. 5.**
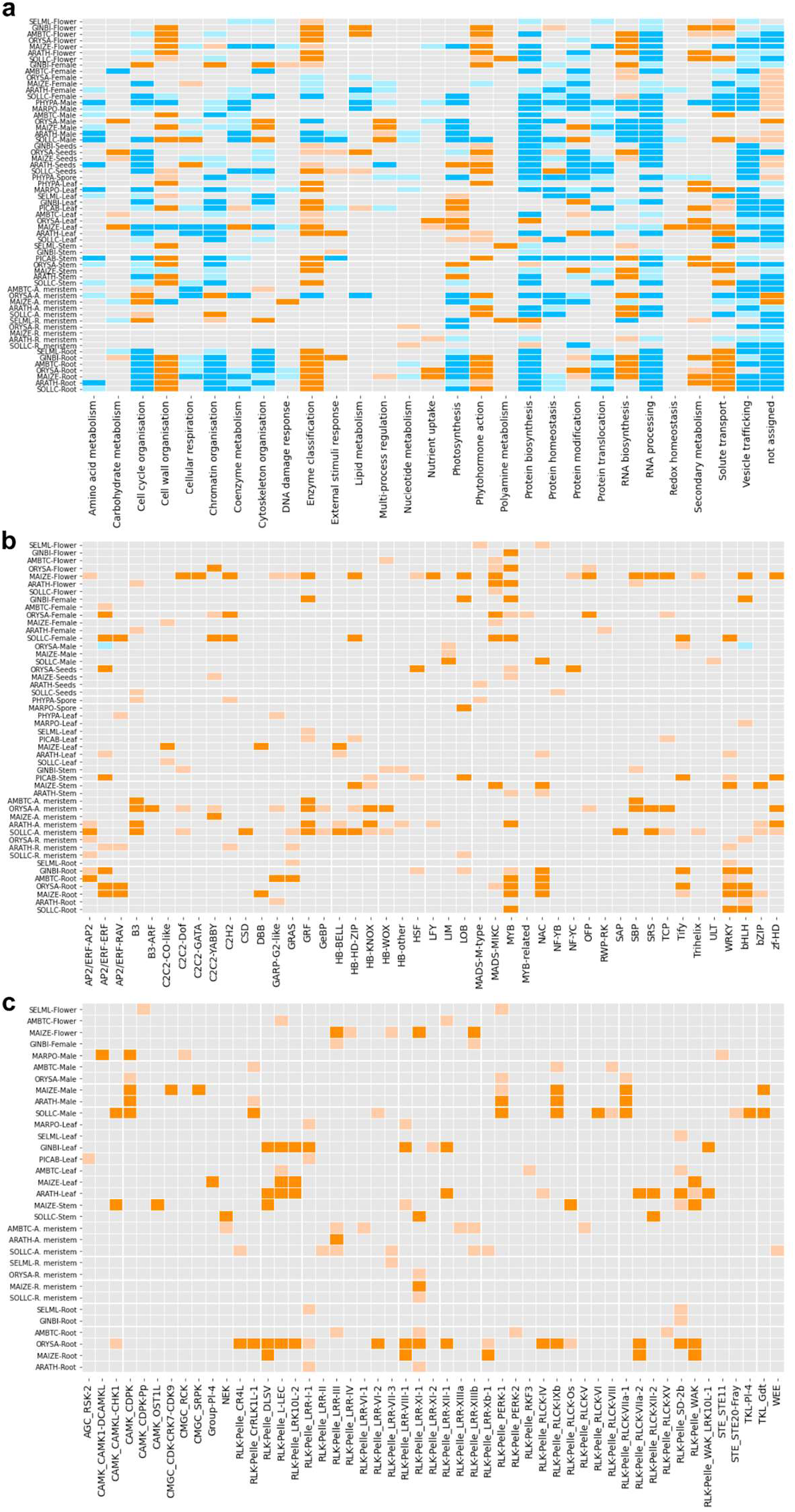
Functional enrichment analysis of the sample-specific transcriptomes. Samples are shown on the y-axis and functions in the x-axis for MapMan bins (a), transcription factors (b), and kinases (c). Orange and blue colors indicate enrichment and depletion, respectively. The intensity of the color is in correlation with the p-value (dark orange/blue: p <0.01, light orange/blue: p<0.05).

**Supplementary Fig. 6:**
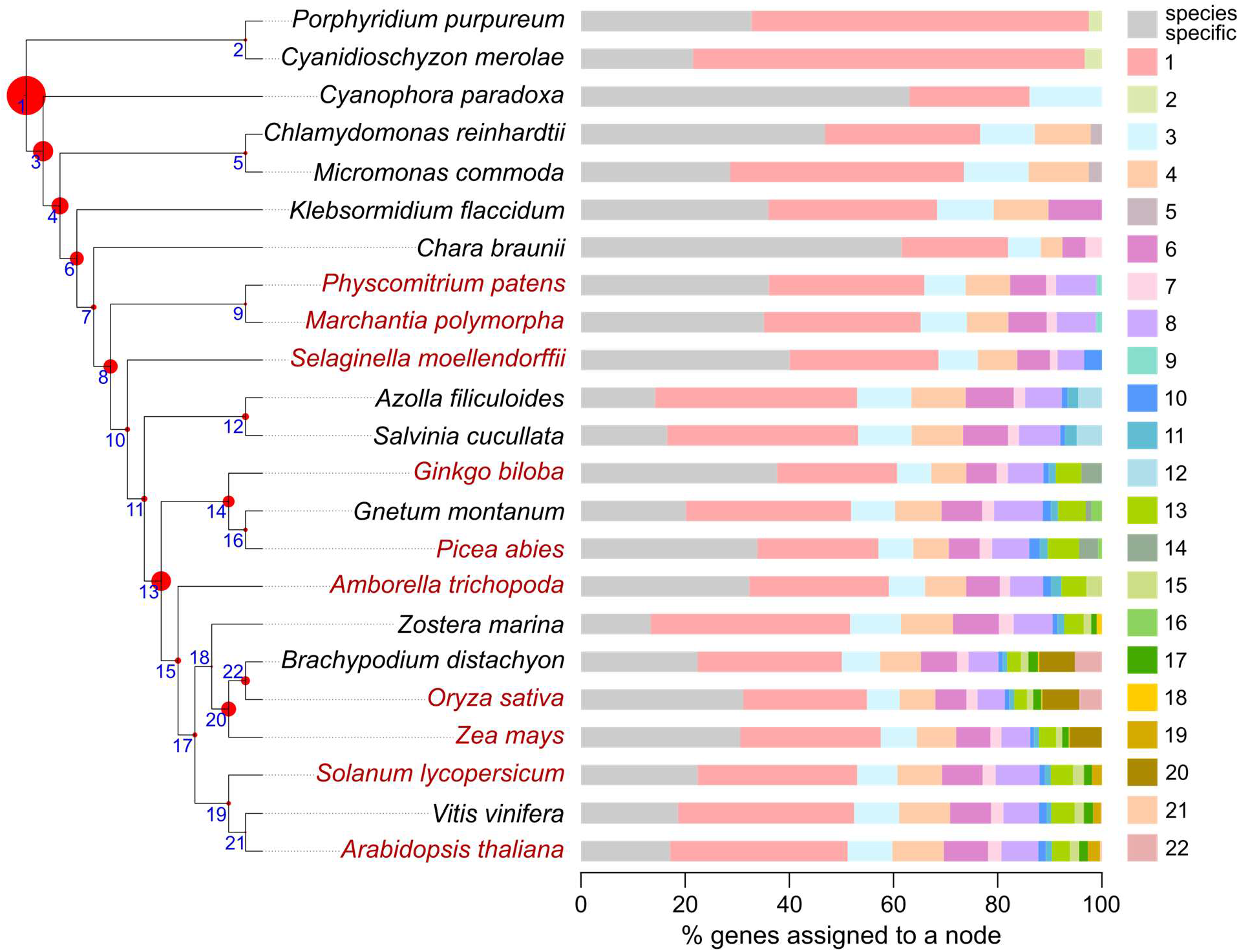
Cladogram of the 23 species included in the analysis. The phylogenetic relationship was based on One Thousand Plant Transcriptomes Initiative, 2019. Species in red are associated with transcriptomic data in this study. Blue numbers in the nodes indicate the node number (e.g., 1: NODE_1). The tree’s red circles show the percentage of orthogroups found in each node (largest and smallest amounts: Node_1 - 24% and NODE_21 - 0.1%). Bars on the right show the percentage of genes per species that are present in each node. The nodes are shown in different colors, as indicated in the right bar.

**Supplementary Fig. 7:**
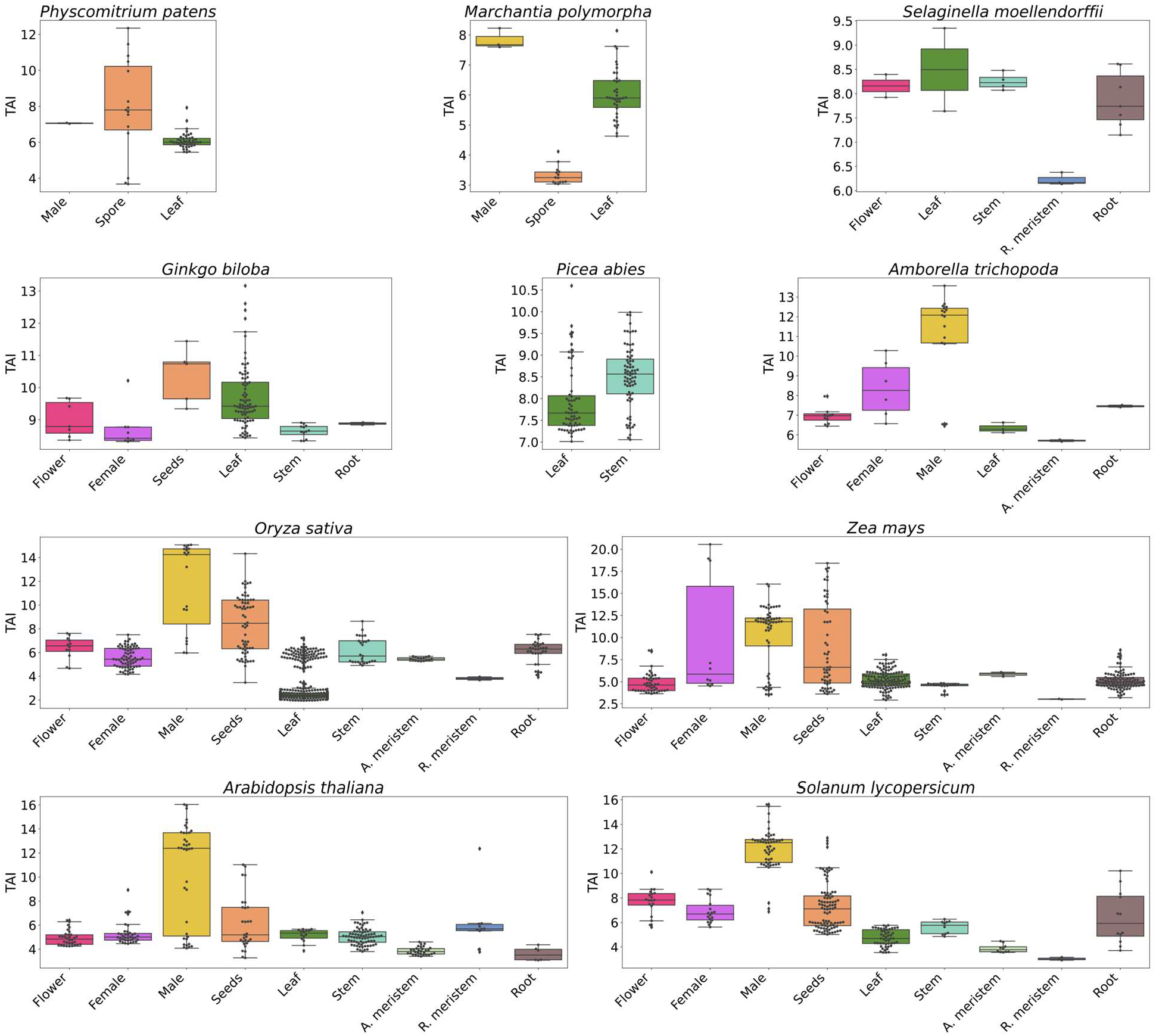
Transcriptomic age index in the ten species.

**Supplementary Fig. 8:**
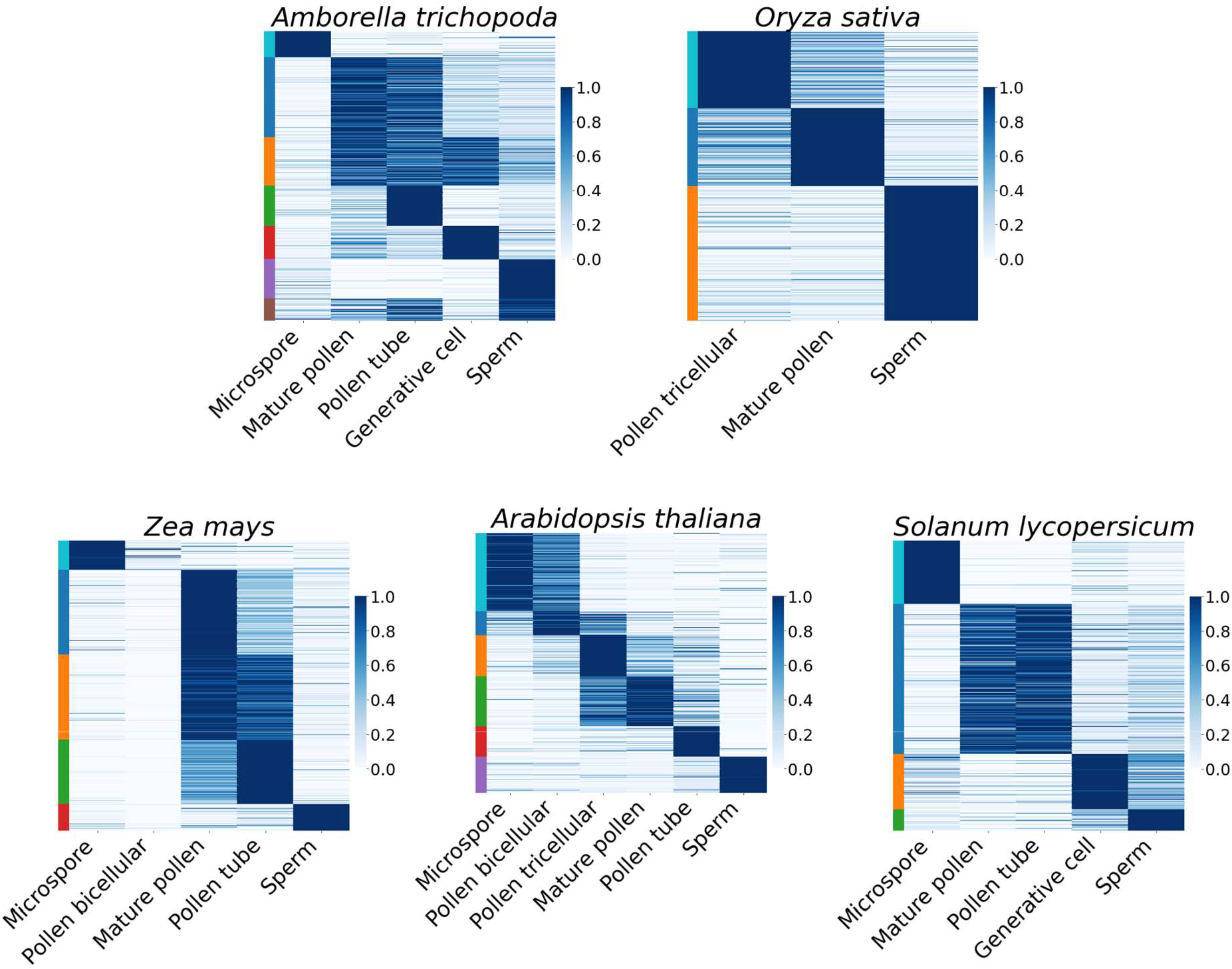
Expression of male developmental stages genes for five species. Genes are in rows, developmental stages in columns. Gene expression is scaled to range from 0-1. Darker color corresponds to a stronger positive correlation. Bars in the left mark the different clusters.

**Supplementary Fig. 9:**
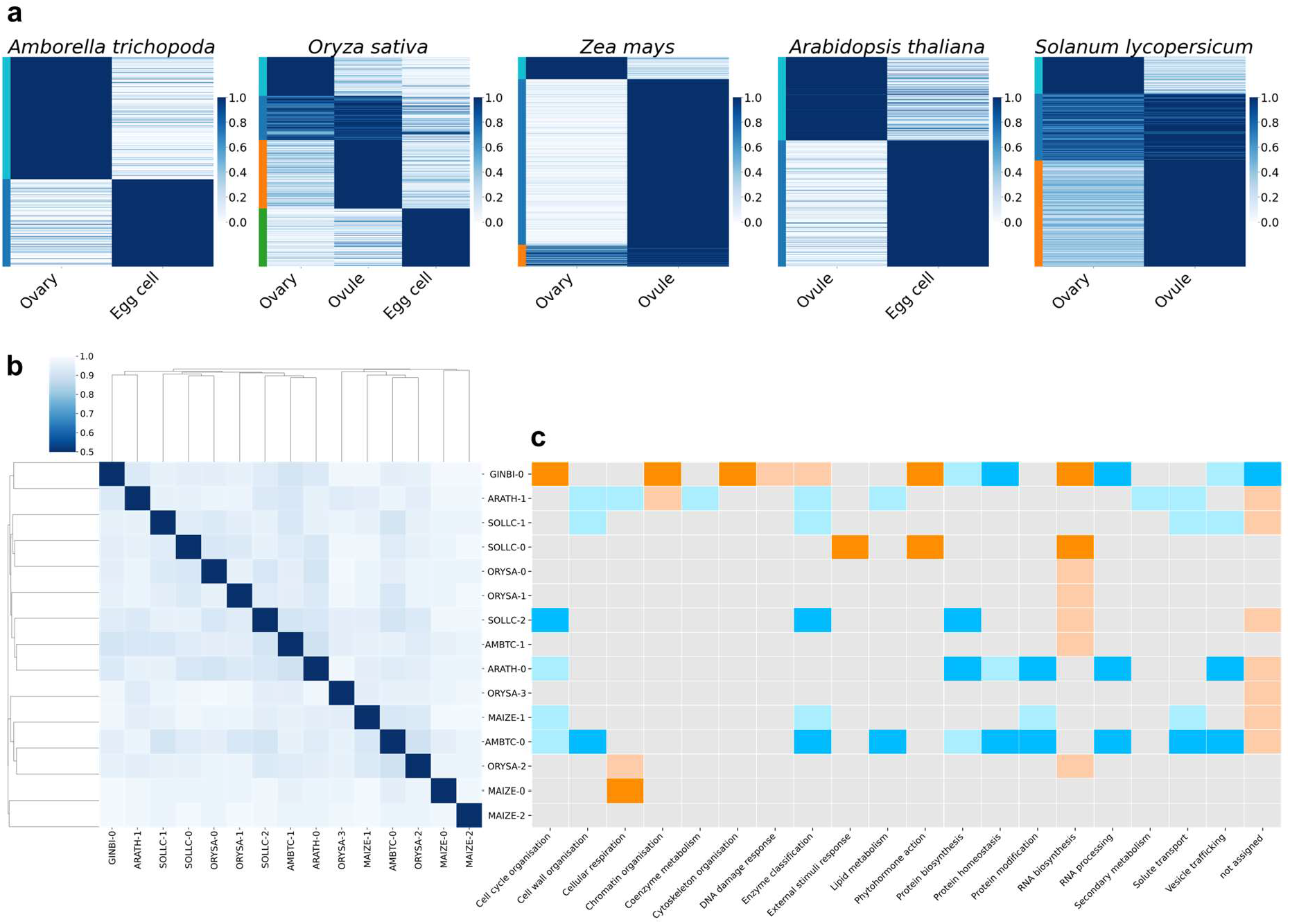
Analysis of the expression profile in different development stages of female organs. Heat map showing the normalized TMP of genes per each development stage for five species. Bars on the left indicate the clusters. **b**, Jaccard distance between the clusters. **c**, Heatmap showing enrichment and depletion of functions. Orange and blue indicate enrichment and depletion, respectively (light colors: p < 0.05, dark colors: p < 0.01).

**Supplementary Fig. 10:**
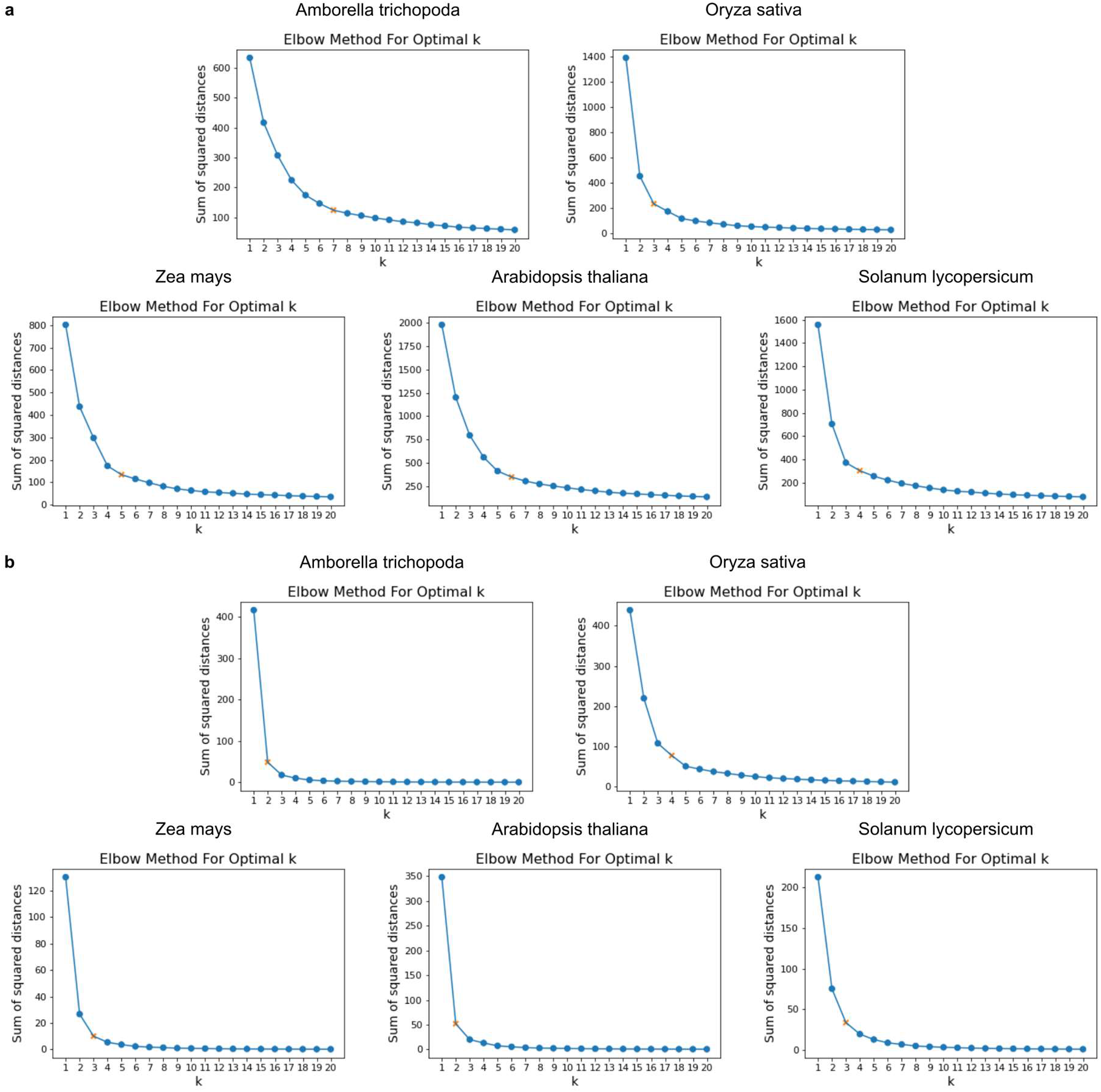
Identifying *k* value with the elbow method. The orange mark indicates the *k* value where the sum of squared distances was less than 80% of the highest value found at *k* =1. **a**, For the male samples, and **b**, for the female samples.

## Supplementary tables

**Supplementary Table 1.** Samples included per each species. The columns in order show: mnemonic of the species, sample ID, original annotation, Organ name, Subsample name, source of the sample, number of fragments that could be pseudoaligned using, percentage of fragments that could be pseudoaligned, tag for the samples that pass/fail Kallisto stats, tag for the samples that pass/fail PCC filter.

**Supplementary Table 2.** The number of expressed and organ-specific expressed genes in the ten species.

**Supplementary Table 3.** Organ-specific genes in the ten land plants. Columns show species mnemonic, sample name, number of genes, gene names.

**Supplementary Table 4**. Organ-specific transcription factors in the ten land plants. Mnemonics indicate the species. The columns indicate transcription factor families.

**Supplementary Table 5.** Organ-specific kinases in the ten land plants. The species are indicated by mnemonics, while the organs are given after the species name. The different families of kinases are given in columns.

**Supplementary Table 6.** List of orthogroups identified in the 23 species included. The columns show the orthogroup name, node in the species tree (Fig. 3a), expression profile, pass/fail filter of the expression profile, list of species (mnemonic). The following columns show the list of genes per species.

**Supplementary Table 7.** Sample-specific gene enrichment for species and for node in the species tree (Fig. 3a). The columns show: species mnemonic, sample name, node of the species tree, p-value, tag (enrichment or depletion).

**Supplementary Table 8.** Gain/loss of gene families. The columns show the sample, node, number of total gains, number of total losses, orthogroups gained, orthogroups lost.

**Supplementary Table 9.** List of enriched functions in gained organ-specific and ubiquitous gene families per each node.

**Supplementary Table 10.** List of male cluster-specific genes. The first column shows the mnemonic of the species. The second, the cluster number. The third to ninth column: the average TPM per each male sample included. The last column: the list of genes of the cluster.

**Supplementary Table 11.** List of female cluster-specific genes. The first column shows the mnemonic of the species. The second, the cluster number. The third to fifth column: the average TPM per each male sample included. The sixth column: the list of genes of the cluster.

**Supplementary Table 12.** Features of male cluster-specific genes. The columns show the mnemonic of the species, gene name, cluster number, if it is co-expressed (Yes/No), transcription factor, or kinase name if reported in the annotation.

**Supplementary Table 13.** Annotation ofthe male cluster-specific genes of A. *thaliana.* The columns show: cluster name, gene, tag for transcription factor (TF) or kinase (KIN), name of the transcription factor or kinase, if it is co-expressed (Y /N), name of the sample that the known mutant affect, mutant, if the gene is involved in pollen, references.

**Supplementary Table 14.** List of species included in this study and the source of their proteomes and CDSs. Columns show: mnemonic, taxon identifier, species name, genome version, and source.

**Supplementary Table 15.** Gene families (first column), Arabidopsis male-specific genes (second column) and Amborella male-specific genes (third column). Gene and family IDs are clickable and will redirect the user to a corresponding page. The fourth column indicates gene families found in common in Arabidopsis and Amborella (intersection), only in Arabidopsis (left) or only in Amborella (right).

